# The SCF/KIT pathway implements self-organised epithelial patterning by cell movement

**DOI:** 10.1101/2020.04.22.055483

**Authors:** Alexandre Chuyen, Charlotte Rulquin, Virginie Thomé, Raphaël Clément, Laurent Kodjabachian, Andrea Pasini

**Author notes:** Co-senior authors (Laurent Kodjabachian, Andrea Pasini).

## Abstract

How individual cell behaviours lead to the emergence of global patterns is poorly understood. In the *Xenopus* embryonic epidermis, multiciliated cells (MCCs) are born in a random pattern within an inner mesenchymal layer, and subsequently intercalate at regular intervals into an outer epithelial layer. Using both experiments and mathematical modelling, we show that this transition from chaotic to ordered distribution relies on mutual repulsion among motile immature MCCs, and affinity towards outer-layer intercellular junctions. Consistently, ARP2/3-mediated actin remodelling is required for MCC pattern emergence. Using multiple functional approaches, we show that the Kit tyrosine kinase receptor, expressed in MCCs, and its ligand Scf, expressed in outer-layer cells, are both required for regular MCC distribution. While Scf behaves as a potent adhesive cue for MCCs, Kit expression is sufficient to confer order to a disordered heterologous cell population. Our work reveals how a single signalling system can implement self-organised large-scale patterning.

**Highlights:**

- Immature multiciliated cells transit from a disordered to an ordered pattern
- The transition is a self-organising process based on repulsive and affinity movements
- ARP2/3-dependent actin remodelling is required for pattern emergence
- The SCF/KIT pathway promotes both repulsion and affinity movements

**eTOC blurb:** In developing *Xenopus* epidermis, immature multiciliated cells (MCCs), initially chaotically distributed within an inner layer, emerge in an orderly pattern among cells of the outer layer. This process involves MCC mutual repulsion and affinity towards outer-layer intercellular junctions. The SCF/KIT signalling pathway promotes both properties to allow regular MCC distribution.

## INTRODUCTION

Orderly cellular patterns have always captured the attention of scientists, and understanding the mechanisms and forces that control their establishment is one of the long-standing aims of developmental biology. In the past decades, classical and molecular genetics approaches have led to the identification of robust Gene Regulatory Networks (GRNs) that generate patterns by concomitantly specifying cellular identity and position within developing embryos (Briscoe and Small, 2015; Davidson, 2010). However, patterns of regular cell distribution can also emerge when the stochastic movements of motile cell populations are partially restricted, for example by mutual repulsion. Such phenomena have been described in rat retina (Galli-Resta et al., 2002), mouse cerebral cortex (Villar-Cervino et al., 2013), zebrafish epidermal pigmented cells (Walderich et al., 2016) and *Drosophila* haemocytes (Davis et al., 2012) but the underlying molecular mechanisms are only starting to be unravelled.

In vertebrates, one striking example of ordered cellular pattern is provided by the embryonic epidermis of the amphibian *Xenopus*, where hundreds of cells carrying multiple motile cilia (Multiciliated Cells or MCCs) are found distributed among mucus-producing goblet cells, according to a regular spacing pattern (Deblandre et al., 1999). The ontogeny of this mucociliary epithelium occurs through a multistep process (Cibois, 2014). At cleavage stages, asymmetric cell divisions partition the embryonic non-neural ectoderm into an outer epithelial and an inner mesenchymal layer. At gastrula stages, the outer layer gives rise to a sealed epithelium containing only goblet cells, while MCCs are born within the inner layer, together with osmoregulatory ionocytes, serotonin-secreting Small Secretory Cells (SSCs) and basal cells. The BMP and Delta/Notch pathways have been shown to control the number of MCCs, ionocytes and SSCs born within the inner layer (Deblandre et al., 1999) (Stubbs et al., 2006) (Hayes et al., 2007) (Quigley et al., 2011) (Cibois et al., 2015). Starting at neurula stages, the MCCs, ionocytes and SSCs migrate apically in separate waves to radially intercalate into the outer layer, where they will complete their differentiation (Cibois, 2014). Strikingly, however, only MCCs display a regular pattern of distribution in the mature epithelium. Although the precise physiological function of *Xenopus* MCCs remains to be addressed, a regular pattern of distribution may be optimal to homogenize fluid flows powered by beating cilia. MCCs always insert into the outer layer individually and exclusively in correspondence of junctions among at least three outer-layer goblet cells (vertices) (Chung et al., 2014; Deblandre et al., 1999; Stubbs et al., 2006). Radial intercalation involves a first phase in which immature MCCs in the inner layer emit apically-polarized protrusions to probe outer-layer vertices, followed by a phase of apical expansion, when their apical aspect actually emerges at the surface of the outer layer and progressively expands in its plane to allow for cilia assembly (Chung et al., 2014; Stubbs et al., 2006)(Sedzinski et al., 2016). While multiple studies have addressed the cellular and molecular mechanisms of radial MCC intercalation (Sirour et al., 2011)(Kim et al., 2012)(Chung et al., 2014)(Werner et al., 2014)(Sedzinski et al., 2016, 2017), how the regularly dispersed pattern of mature MCCs is generated remains to be resolved. It has been suggested that MCC pattern formation may result from an interplay between the mechanisms that define the numbers of committed immature MCCs and the number and distribution of outer-layer vertices available for intercalation (Deblandre et al., 1999; Stubbs et al., 2006; Werner and Mitchell, 2012). While these two parameters are indisputably involved, it is unclear whether they are sufficient to explain pattern emergence. Here, we hypothesize that MCC movement may be another key ingredient, which has never been evaluated. Using transient transgenesis, we could record the movements of immature MCCs in the inner layer, and concluded that MCC spacing pattern was largely established prior to intercalation. Using mathematical modelling, we found that mutual repulsion among MCCs and affinity for outer-layer intercellular junctions are sufficient to account for the final dispersed pattern. These two behaviours were readily apparent from our live recordings. Confirming the importance of MCC movements, inhibition of ARP2/3-mediated actin remodelling impaired lamellipodia-like protrusions in MCCs, homotypic MCC repulsion and MCC spacing pattern, without affecting intercalation. We also uncover a hitherto unknown role of the signalling pathway dependent on the interaction between the transmembrane tyrosine kinase receptor Kit, expressed in MCCs, and its ligand Scf, expressed in outer-layer cells, for MCC pattern emergence. The SCF/KIT signalling pathway has a well-established role in controlling actin cytoskeleton and cell motility in a variety of vertebrate systems (Lennartsson and Ronnstrand, 2012; Meininger et al., 1992; Runyan et al., 2006; Samayawardhena et al., 2007; Wehrle-Haller et al., 2001; Zuo et al., 2016) and KIT stimulation with SCF was found to result in the phosphorylation and activation of ARP2/3 (Mani et al., 2009). Using a variety of functional assays, we established the capacity of the Scf/Kit system to promote both the repulsive and adhesive interactions predicted *in silico*, and thus identified for the first time a major regulator of the *Xenopus* MCC patterning process. This work illustrates how a large cell population can self-organise to adopt a regular spacing pattern, through semi-random movements constrained by a balance between repulsive and attractive cues.

## RESULTS

### Emergence of MCC ordered pattern precedes radial intercalation

Fluorescent In Situ Hybridization (FISH) of *Xenopus* embryos with the MCC marker *α-tubulin* (*α-tub*) revealed that at stage 14 (early neural plate stage), immature MCCs were irregularly distributed within the non-neural ectoderm inner layer (Figure 1A) and very often in direct contact with each other (Figure 1B). Co-staining with an antibody against the tight junction protein ZO-1, to mark outer-layer apical tight junctions, also showed that at this stage *α-tub*-positive MCCs were not positioned in correspondence of outer-layer vertices (Figure 1A, 1C). In the course of development, however, *α-tub*-positive MCCs were increasingly often found as separate cells (Figure 1A, 1B), and progressively localized each in correspondence of an individual vertex (Figure 1A, 1C), into which they finally inserted, thus progressively taking up an apparently more orderly pattern (Figure 1A). To quantify the progression from an irregular to a regular distribution, we developed an order index based on Delaunay triangulation of the centroids of *α-tub*-positive cells. Such an index is inversely proportional to the variation of distances among centroids and would reach its highest value when MCC centroids are perfectly ordered in staggered rows. As shown in Figure 1D, the order index of the MCC population progressively increased from stage 14 through stage 25, when MCCs are intercalated into the outer layer.

**Figure 1.**
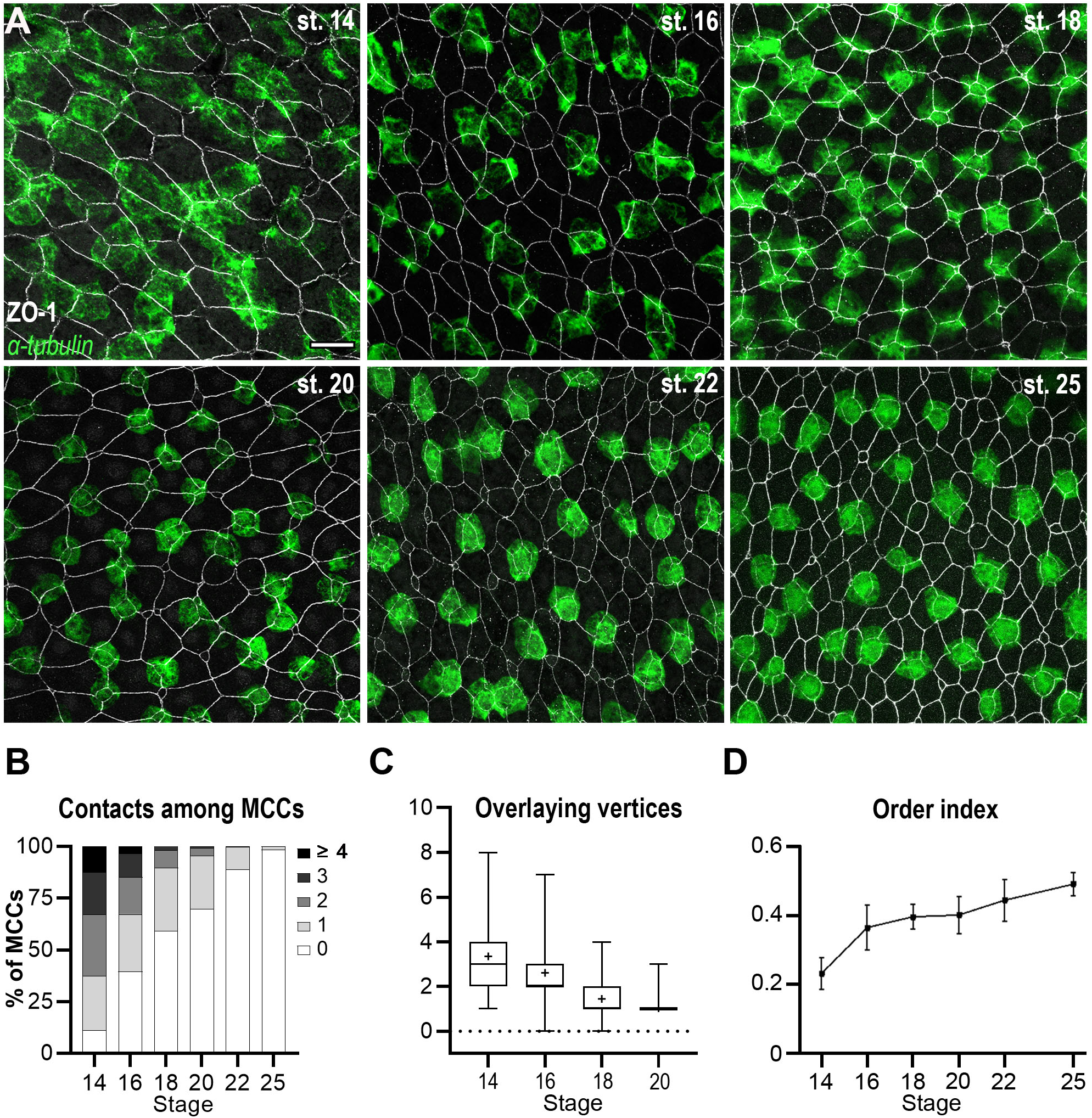
The orderly pattern of mature MCCs results from progressive changes in cell distribution. **A:** Fluorescent *In Situ* Hybridization (FISH) was used to reveal *α-tubulin*-positive immature MCCs (green), combined to immunofluorescence against the tight junction protein ZO-1 to reveal outer-layer intercellular junctions (white). In the course of development, MCCs were increasingly found as isolated, regularly-shaped cells, each localised in correspondence of a single outer-layer vertex. Scale bar is 25μm. **B:** Graph showing the percentage of *α-tub*-positive cells directly in contact with varying numbers of other *α-tub*-positive cells (gray-shaded from 0 to to ≥4). **C:** Graph showing the number of ZO-1-positive outer-layer vertices superimposable onto the planar projection of each *α-tub*-positive MCC. **D:** Graph showing the order index calculated from Delaunay tessellation of the centroids of *α-tub*-positive cells. The order index increased during development, reflecting the progressive transition from a chaotic to an orderly MCC distribution.

The apparent increase in order may result non-exclusively from selective death or cell fate reversal of misplaced MCCs, or from MCC movements. Fluorescent detection of the apoptotic executioner Caspase 3/7 activity as a proxy of apoptosis failed to reveal dying cells in the ectoderm of control embryos from stage 15 to stage 18, as opposed to cycloheximide-treated embryos (Trindade et al., 2003) (Figure S1A and movie S1). To evaluate the possible contribution of cell-fate reversal, we performed *α-tub* FISH in embryos from a transgenic line expressing RFP under the control of the MCC-specific α-tub promoter (Stubbs et al., 2006). Due to the long RFP half-life (26h), any immature MCC switching its identity to that of a different inner layer cell type should retain RFP fluorescence while losing endogenous *α-tub* transcripts. Such cases were never observed, ruling out MCC fate reversal during normal development (Figure S1B, S1C). In contrast, we noticed that robust *α-tub* signal was occasionally detected in RFP-negative cells, compatible with events of asynchronous MCC specification (Figure S1B, S1C).

The lack of MCC death or fate reversal led us to favour the hypothesis that cell motility might play a major role in the progressive emergence of a regular pattern of MCC distribution.

### Time-lapse video microscopy reveals MCC motility

To evaluate whether MCCs are motile, we injected embryos with constructs carrying the fluorescent markers GFP or LifeActGFP (Belin et al., 2014) under the control of the α-tub promoter, and followed the behaviour of labelled MCCs in living embryos. Despite the high degree of mosaicism, which restricted transgene expression to a small percentage of MCCs, fluorescence levels were sufficient to allow live recording starting from stage 14 and for a duration of up to 8 hours. Spinning-disk confocal video microscopy showed that GFP-expressing MCCs indeed moved actively within the plane of the inner layer until they stopped in correspondence of vertices where they began to intercalate (Figure 2A and movie 1). As the F-actin-binding reporter LifeActGFP allowed for much better visualisation of the whole cell shape and of cellular protrusions, we exploited it for most of our subsequent experiments. We thus found that LifeActGFP-expressing MCCs extended and retracted actin-rich lamellipodia-like processes within the ectoderm inner layer and progressively changed their shape from flat mesenchymal to more compact and pyramidal (Figure 2B and movie 2). These observations suggested that cell motility may indeed be important to establish a regular MCC pattern.

**Figure 2.**
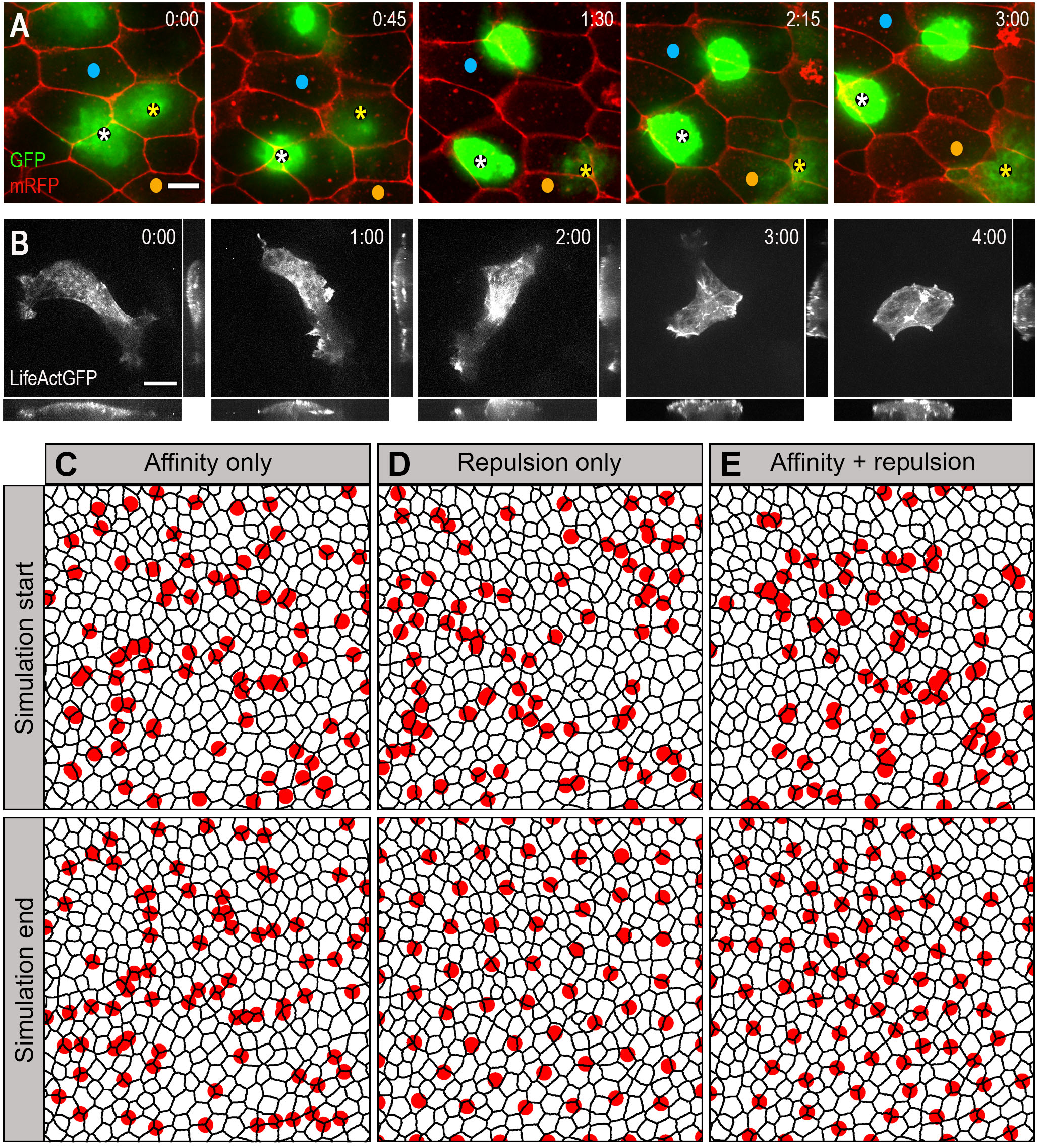
*In silico* modelling of MCC behaviour reveals that mutual repulsion and affinity for outer-layer junctions are required for proper pattern establishment. **A:** Spinning disk confocal video microscopy was used to track α-tub::GFP transgenic MCCs within the inner epidermal layer. Still frames from movie 1 are shown. Asterisks point two MCCs that moved under mRFP-injected outer-layer cells. Blue and orange dots serve as landmarks to appreciate MCC mobility. **B:** Still frames from movie 2. α-tub::LifeActGFP revealed an intense actin-based protrusive activity that progressively shifted from the basolateral to the apical aspect of the cell. **C-E:** Frames from simulation movies 3, 4 and 5, respectively. The black meshwork represents outer-layer intercellular junctions, and red circles the underlying MCCs. The upper row shows the initial random distribution of MCCs, the bottom row shows the final patterns obtained under the different regimes. **C:** In a regime where MCC affinity for outer-layer junctions dominated, MCCs localised at vertices, but failed to disperse properly. **D:** In a regime where MCC mutual repulsion dominated, MCCs were regularly distributed, but did not localise at vertices. **E:** In a regime where mutual repulsion and affinity for outer-layer junctions were balanced, MCCs were dispersed and positioned at vertices.

### A cell motility-based mathematical model for MCC pattern establishment

We next built an energy-based mathematical model to identify the minimal ingredients necessary to recapitulate the observed self-organisation of motile MCCs. In this simplified model, MCCs can move in a bi-dimensional plane against a network representing outer-layer junctions (see STAR methods). As homotypic repulsion is often required for homogeneous dispersion of motile cellular populations (Davis et al., 2012; Villar-Cervino et al., 2013), we introduced a first term related to MCC mutual repulsion, such that proximity to another MCC increased energy (see STAR methods). In addition, we reasoned that homing of MCCs to outer-layer vertices must imply a marked affinity for epithelial junctions. Thus, we introduced a second term related to affinity, such that the overlap of MCCs with junctions decreased energy (see STAR methods). The model also included a steric constraint in the form of a hard-core potential, to prevent complete overlap between MCCs. Starting from a random spatial distribution of MCCs, a standard Monte-Carlo scheme was used to allow system evolution. At each step, any given MCC was assigned a random, diffusion-like prospective displacement. A displacement was systematically accepted if it lowered the system total energy. If on the contrary it increased system energy, its acceptance probability decayed exponentially with energy increase (see STAR methods). This eventually led the system towards equilibrium configurations. In an affinity-dominated regime (i.e. when the contribution of mutual repulsion is negligible), MCCs rapidly re-located at nearby vertices, where they could maximise their interaction with outer-layer junctions and minimize their energy. However, this occurred regardless of the position of other MCCs, so that no overall regularity of distribution was observed (Figure 2C and movie 3). In contrast, in a repulsion-dominated regime, MCCs repelled each other and distributed according to a regular pattern that maximized their mutual distance, regardless of the position of outer-layer vertices (Figure 2D and movie 4). Finally, in an intermediate regime in which the repulsion and affinity terms were commensurate, a trade-off situation was observed, in which MCCs both located at outer-layer vertices and dispersed homogeneously (Figure 2E and movie 5; also see STAR methods). Thus, self-organised MCC distribution can be captured in a simple mathematical model where movements are constrained by homotypic repulsion and affinity for outer-layer junctions.

### Mutual repulsion and affinity for outer-layer junctions underlie MCC pattern establishment

We next explored whether MCC mutual repulsion and affinity for outer-layer junctions predicted to be important in our *in silico* model were indeed detectable *in vivo*. In cases where low mosaicism allowed for visualisation of several neighbouring LifeActGFP-expressing MCCs we found that, whenever in the course of their movements two such cells came into contact, they withdrew from each other (Figure 3A and S2A and movie 6). Overall, the average time of contact between two LifeActGFP-expressing cells was calculated at 12.11+/-18.96 minutes (average +/- SD) (Figure 3F). On the other hand, imaging of embryos simultaneously injected with α-tub::LifeActGFP and with a synthetic mRNA coding for a membrane-localized form of RFP to label outer-layer cells revealed that, while in the inner layer, most MCCs preferentially moved along the junctions among outer-layer cells (92.55% of 94 analysed MCCs closely followed outer-layer junctions over >2hrs) (Figure 3B, 3G and movie 7). These observations confirmed that homotypic repulsion and affinity for outer-layer junctions are likely important for the emergence of a regular MCC pattern.

**Figure 3.**
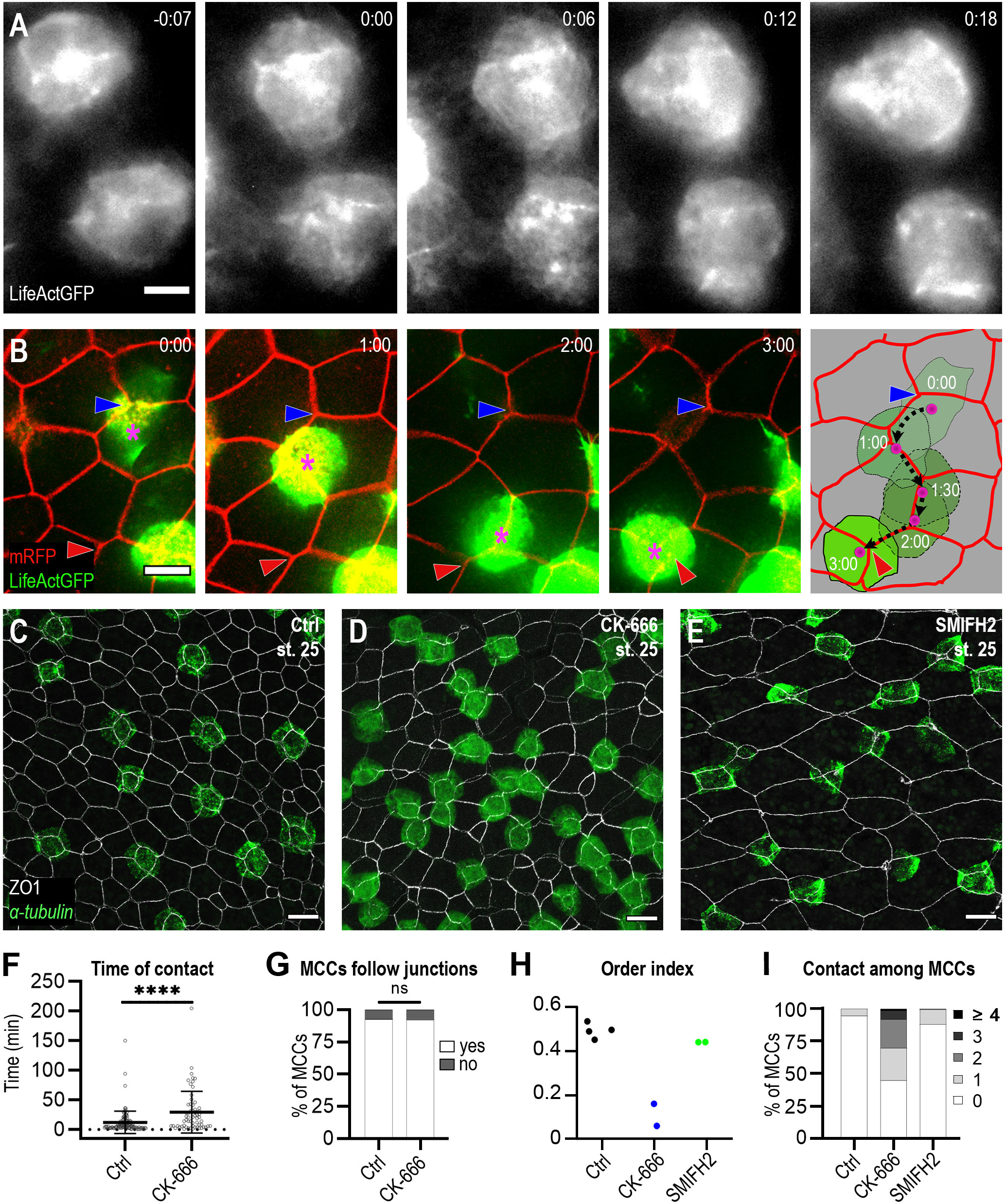
MCCs display both behaviours of mutual repulsion and affinity for outer-layer junctions. **A:** Frames from movie 6. α-tub::LifeActGFP labelling revealed that whenever MCCs came in contact via actin-based lamellipodia, these subsequently retracted. **B:** Frames from movie 7 showing an α-tub::LifeActGFP-labelled MCC (magenta asterisk) moving along the intercellular junctions among overlying outer-layer cells labelled by mRFP. The blue and the red arrowheads highlight the initial and final position of the labelled MCC. The last panel recapitulated the positions of the observed MCC over 3h. The black dotted line shows the displacement of the MCC centroid. **C-E:** Embryos were treated from stage 12 with 1% DMSO (C), 100μM ARP2/3 inhibitor CK-666 (D), and 50μM FORMIN inhibitor SMIFH2 (E) and analysed at stage 25 by *α-tub* FISH and ZO-1 IF to examine MCC patterning. **D:** Following CK-666 treatment, MCCs intercalated into the outer layer but were irregularly distributed. **E:** Treatment with SMIFH2 led to a normal MCC patterning but impaired apical insertion and expansion. In all cases, scale bar is 20μm. **F:** Graph showing the duration of MCC mutual contacts in control and CK-666 treated embryos.**** p-value < 0.0001 on a student t-test. **G:** Graph showing the proportion of MCCs that followed outer-layer junctions over a period of 2h. **H:** Graph showing MCC order index at stage 25 in CK-666-treated and SMIFH2-treated embryos vs control embryos. **I:** Graph showing the distribution of the number of contacts among MCCs at stage 25 in control, CK-666- and SMIFH2-treated embryos.

The actin cytoskeleton appeared to be involved in both MCC mutual repulsion and in their affinity for outer-layer junctions. MCCs often established transient contacts with each other via LifeActGFP-enriched lamellipodia-like structures that were subsequently retracted (Figure 3A and S2A) and LifeActGFP-rich cellular protrusions were apically extended by MCCs as they crawled along junctions among outer-layer cells (Figure S2B and movie S2).

As a first step towards dissecting the role of the actin cytoskeleton in MCC patterning, we took advantage of well-characterised pharmacological inhibitors of the two main pathways of actin polymerisation: CK-666, an inhibitor of the actin-branching nucleator ARP2/3 (Hetrick et al., 2013) and SMIFH2, an inhibitor of the actin-bundling factor FORMIN (Rizvi et al., 2009). FISH for *α-tub* coupled with ZO-1 IF on embryos treated with 100μM CK-666 from stage 12 and fixed at stage 25 showed that most MCCs had properly emerged at the outer-layer apical surface (Figure S2C), but were irregularly distributed and often in direct contact with one another (Figure 3D, 3H, 3I). Consistent with this observation, spinning-disk confocal video microscopy of α-tub::LifeActGFP- and mRFP-injected embryos treated with CK-666 showed a decrease in mutual repulsion among MCCs (average time of contact +/- SD: 48.54+/-49.10 min) (Figure 3F) and formed small clusters (movie S3). On the other hand, in CK-666-treated embryos, MCCs still moved preferentially along outer-layer junctions (92.31% of 26 analysed MCCs) (Figure 3G). An apparent increase in the occurrence of spiky filopodia-like structures over broader, lamellipodia-like ones (Figure S2D, S2D’ and movie S3), reminiscent of what has been described in ArpC3^-/-^ murine fibroblast (Suraneni et al., 2015) was also observed. Interestingly, a fusion protein between the p41 subunit of the actin-branching nucleator ARP2/3 and EGFP, specifically expressed in MCCs by means of the α-tub promoter together with LifeActRFP, accumulated just behind the leading edge of the LifeActRFP-rich protrusions and was quickly relocated when these collapsed and reformed at different cellular locations following MCC contacts (Figure S2E and movie S4). In contrast to what we observed following CK-666 treatment, in embryos treated from stage 12 with the FORMIN inhibitor SMIFH2 at 50μM, MCCs were properly dispersed and positioned at vertices, but their apical emergence at stage 25 was delayed (Figure 3E, 3H, 3I and Figure S2C), in agreement with published data (Sedzinski et al., 2016). Our results thus reveal the importance of actin cytoskeleton-mediated cell movements in MCC patterning and uncover a possible functional dichotomy between actin branching, required for MCC mutual repulsion and regular distribution, and actin bundling, involved in the actual intercalation and apical emergence of MCCs.

### The Receptor Tyrosine Kinase Kit is expressed in MCCs, its ligand Scf in outer-layer cells

Next, we searched for a molecular mechanism that could sustain self-organised MCC pattern emergence. We reasoned that both homotypic repulsion and affinity towards outer-layer junctions must involve cell surface molecules. Our attention was drawn to the Receptor Tyrosine Kinase (RTK) Kit, which has been identified as a transcriptional target of the MCC-inducing factor Multicilin/Mcidas in *X. laevis* epidermis (Kim et al., 2018; Ma et al., 2014; Stubbs et al., 2012). In addition, its transcripts have been detected by single-cell transcriptomics in *X. tropicalis* immature and mature MCCs (Briggs et al., 2018) (https://kleintools.hms.harvard.edu/tools/currentDatasetsList_xenopus_v2.html). As the signalling pathway dependent on KIT and its ligand SCF (Stem Cell Factor, also known as KITLG, Mast Cell Growth Factor or Steel Factor) is known to control actin cytoskeleton and cell motility, we hypothesized that it might also be involved in controlling the cellular behaviours that underlie the regular pattern of *Xenopus* MCCs. Whole-mount ISH with an RNA probe recognizing transcripts of both *kit-L* and *kit-S* homeologs detected expression by scattered cells in the non-neural ectoderm starting from early gastrula (stage 10) and persisting through later developmental stages (stage 26) (Figure S3A). FISH on sectioned embryos revealed that in the developing epidermis *kit* transcripts were exclusively co-expressed with the MCC markers *foxj1* and *α-tubulin* (Figure 4A). Conversely, expression throughout the developing non-neural ectoderm has been reported for *scf-L* and, at much lower levels, for *scf-S* (Goldman et al., 2006; Martin and Harland, 2004) (Figure S3B). RT-PCR on total mRNA from animal caps confirmed the expression of *scf-L* from stage 12 to 22, but failed to detect *scf-S* (Figure S3C). FISH revealed that *scf-L* transcripts were prevalently localised in the outer layer, with much lower levels of expression in the inner layer (Figure S3B’). In particular, *scf-L* expression was rarely detected and only at very low levels in *kit*-positive MCCs (Figure 4B). Accordingly, single-cell transcriptomics revealed that in *X. tropicalis* non-neural ectoderm, expression of *scf* is highest in goblet cells and excluded from MCCs (Briggs et al., 2018) (https://kleintools.hms.harvard.edu/tools/viewTree.html?treedata/final_180213) In many tetrapods, SCF is synthesised as a transmembrane molecule but alternative splicing of the exon 6 during mRNA maturation can remove a major serine-protease cleavage site. As a result, the exon 6-containing isoform is cleaved to release a soluble peptide, while the exon 6-spliced isoform can only be cleaved at much slower rates at other sites, and thus has a longer half-life as a cell membrane-associated molecule (Brannan et al., 1991; Flanagan et al., 1991; Lennartsson and Ronnstrand, 2012). Both soluble and membrane-anchored isoforms of SCF are able to activate the KIT receptor, albeit with different kinetics and biological outcomes (Gommerman et al., 2000; Lennartsson and Ronnstrand, 2012; Miyazawa et al., 1995; Tabone-Eglinger et al., 2014; Tabone-Eglinger et al., 2012). RT-PCR with primers encompassing exon 6 showed that only the exon 6-containing isoform of *scf-L* was detected in developing *Xenopus* ectoderm (Figure S3C). However, the amino-acid stretches coded by exon 6 in Scf-L and Scf-S have very low similarity to mammalian exon 6 and do not display obvious consensus sites for proteolytic cleavage (Hultman et al., 2007). To address the subcellular localisation of *Xenopus* Scf, we generated a construct coding for a C-terminal EGFP-tagged version of Scf-L (Figure 4C) and expressed it in the embryonic ectoderm alone or together with an RFP-tagged version of the tight junction protein ZO-1 (Higashi et al., 2016). Scf-GFP chimeric protein was primarily detected in the lateral plasma membrane of outer ectodermal cells, just basal to ZO1-RFP (Figure 4D, 4D’). However, Western blotting of lysates of Scf-GFP-injected animal caps with an anti-GFP antibody revealed the presence of a peptide compatible with cleavage at exon 6, together with the uncleaved full-length protein (Figure 4E). We thus assume that both membrane-anchored and soluble forms of Scf-L may be present in the developing *Xenopus* ectoderm.

**Figure 4.**
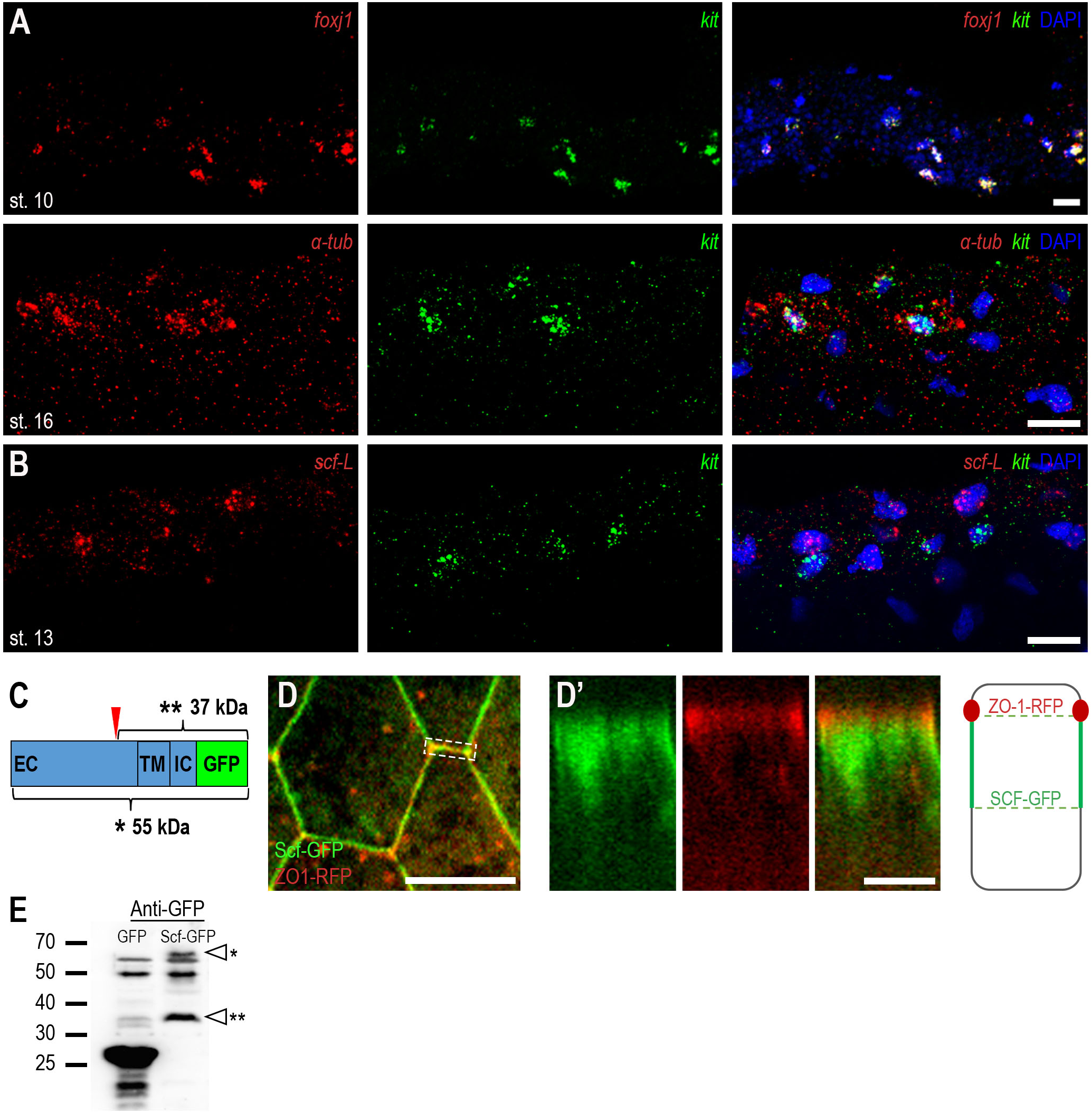
Expression of transcripts encoding Kit and its ligand Scf in the developing *Xenopus* epidermis. **A:** Double FISH on sectioned embryos with probes against *kit* (green) and the MCC markers *foxJ1* or *α-tubulin* (red) revealed that *kit* was only expressed in MCCs. **B:** Double FISH on sectioned embryos showed that *scf-L* (red) expression was prominent in outer-layer cells and virtually undetectable in *kit*-positive MCCs (green). In **A** and **B**, DAPI (blue) highlights the nuclei. **C:** Schematic drawing of the Scf-GFP chimera, showing the extracellular (EC), transmembrane (TM) and intracellular (IC) domains of Scf-L (light blue), the C-terminal EGFP tag (green), the possible protease cleavage site in exon 6 (red arrowhead), as well as the sizes of the full-length protein and the membrane-retained cleavage product. **D:** Top view of live outer-layer cells from an embryo injected with mRNAs coding for the Scf-GFP (green) and the RFP-ZO-1 (red) fusion proteins. GFP fluorescence was localised at the membrane. Scale bar is 20μm. **D’:** Orthogonal view of the intercellular junction within the dashed white box in **D** shows that the Scf-GFP chimera was mainly localised to the basolateral membrane, immediately basal to ZO-1. Scale bar is 5μm. **E:** Lysates from animal caps injected with mRNA coding for either GFP or Scf-GFP were subjected to Western blotting with an anti-GFP antibody. Single and double asterisks point respectively to the 55kDa full-length Scf-GFP and to the 37kDa band corresponding to the membrane-retained product of proteolytic cleavage at exon 6.

### Perturbations of the Scf/Kit signal disrupt MCC patterning

As a first step towards addressing the possible role of Kit signalling in MCC pattern establishment, we took advantage of the KIT tyrosine-kinase pharmacological inhibitor, axitinib (Hu-Lowe et al., 2008). Spinning-disk confocal video microscopy of α-tub::LifeActGFP- and mRFP-injected embryos treated with 50μM axitinib from stage 12 showed that MCCs displayed strongly decreased mutual repulsion (Figure 5A and movie 8) with a time of contact of 59.20+/- 58.15min (average+/-SD) (Figure 5C), but maintained crawling along outer-layer junctions (91.07% of 56 analysed MCCs) (Figure 5C’). An *α-tub* FISH/ZO-1 IF time course on axitinib-treated embryos confirmed that MCCs were largely able to intercalate but did not disperse properly (Figure 5E, 5G-I). As an alternative method to impair Kit activity, we injected a dominant-negative form lacking the TK domain of the receptor (Goldman et al., 2006). Consistently, dn-Kit also caused severe disruption of MCC spacing, although apical emergence and ciliogenesis were largely preserved (Figure S4 and movie S5). Only few Kit-deficient MCCs, often still in direct contact to already inserted ones, were found trapped within the internal layer at stage 25 (Figure S4D’).

**Figure 5.**
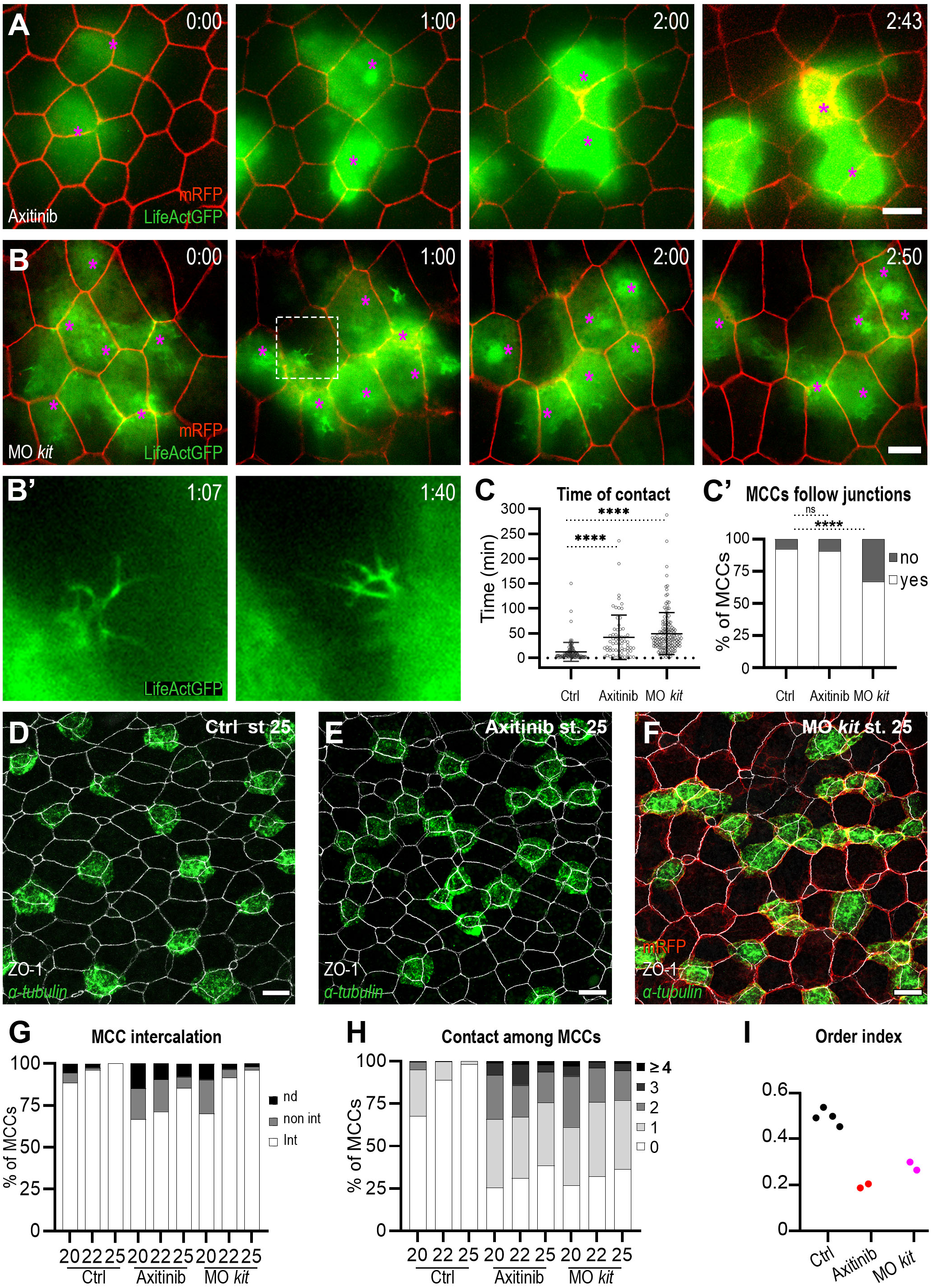
Inhibition of Kit activity disrupts MCC patterning. **A:** Frames from movie 8. In 50μM axitinib-treated embryos, LifeActGFP-positive MCCs (green, magenta asterisk) failed to properly separate from each other. **B:** Frames from movie 9. In MO-ATG-kit-L+S-injected embryos, LifeActGFP-positive MCCs (green, magenta asterisk) failed to properly separate from each other. In **A** and **B**, mRFP (red) labelled outer-layer cell membranes. **B’:** A close-up of the boxed area in **B**, showing highly-branched filopodia-like structures. **C:** Quantification of MCC mutual contact times in control, axitinib-treated and MO-ATG-kit-L+S-injected embryos.**** p-value < 0.0001 on a student t-test. **C’:** Graph showing the proportion of MCCs that followed outer-layer junctions over a period of 2h in control, axitinib-treated, and *kit* MO-injected embryos. **D-F:** *α-tub* FISH (green) and ZO-1 IF (white) showed that compared to control embryos (**D**) MCCs were irregularly distributed in axitinib-treated (**E**) and *kit* MO-injected (**F**) embryos at stage 25. In **F**, mRFP mRNA (red) was used as *kit* MO tracer. In all cases, scale bar is 20μm. **G:** Graph showing the percentage of MCCs intercalated into the outer layer at stages 20, 22 and 25. Intercalation was scored positive when MCCs displayed ZO-1 junctions. Kit inhibition delayed but did not prevent intercalation. **H:** Graph showing the distribution of the number of contacts among MCCs at stage 25 in control, axitinib-treated and *kit* MO–injected embryos The progressive dispersion of *α-tub*-positive MCCs is disrupted in Kit-deficient conditions. **I:** Graph showing MCC order index at stage 25 in axitinib-treated and *kit* MO-injected embryos vs control embryos.

To further confirm the involvement of Kit in MCC pattern establishment, we designed morpholino-modified antisense oligonucleotides (MOs) to block the translation of *kit-L* and *kit-S* mRNAs (MO-ATG-kit-L and MO-ATG-kit-S), or their splicing at the exon2-intron2 junction (MO-splice-kit-L and MO-splice-kit-S) (Figure S5A, S5B). After validation (Figure S5C, S5D), MOs were injected in the two animal ventral blastomeres of 8-cell embryos as cocktails of either MO-ATG-kit-L+S (10ng/blastomere) or MO-splice-kit-L+S (15ng/blastomere). Similar to axitinib-treated and dn-Kit-injected embryos, *α-tub* FISH and ZO-1 IF at stage 25 showed that *kit* MO-injected MCCs were properly intercalated but irregularly distributed and present in small clusters (Figure 5F-I). Co-injection of *kit* MOs with an mRNA coding for full-length Kit-S and carrying seven silent mutations rescued MCC dispersion, thus confirming that this phenotype could be specifically attributed to Kit inactivation (Figure S5E, S5E’). As splicing- and translation-blocking *kit* MOs yielded the same results (Figure S5F, S5F’), all subsequent experiments were performed with the latter ones. Thus, spinning-disk confocal video microscopy of embryos injected with α-tub::LifeActGFP and 10ng/blastomere *kit* MO revealed that MCCs failed to separate from each other (average time of contact +/- SD: 58.48+/-53.31 min) and were compromised in their ability to follow outer-layer intercellular junctions (67.14% of 70 analysed MCCs followed junctions) (Figure 5B-C’ and movie 9). Remarkably, *kit* MO-injected MCCs appeared to emit more filopodia-like than lamellipodia-like structures, similar to CK-666 treatment (Figure 5B’ and movie 9).

As Kit function is likely to reflect interaction with its cognate ligand Scf, we then proceeded to address the effect of Scf knockdown on MCC pattern. MOs were designed to block the translation of both *scf-L* and *scf-S* mRNAs (MO-ATG-scf-L/S) (Goldman et al., 2006), or the exon2-intron2 splicing of the *scf-L* mRNA (MO-splice-scf-L) (Figure S6A, S6B). After validation (Figure S6C, S6D), 10ng/blastomere of MO-ATG-scf-L/S was injected in epidermal precursor blastomeres at 8-cell stage. *α-tub* FISH and ZO-1 IF at stage 25 showed defects in MCC dispersion (Figure 6A-D), while insertion into the outer layer appeared largely normal (Figure S6F). Injection of 10ng/blastomere of MO-splice-scf-L led to the same phenotype as MO-ATG-scf-L/S (Figure S6E, S6E’) and this latter MO was used for all subsequent experiments. Spinning-disk confocal video microscopy of embryos injected with α-tub::LifeActGFP and Scf MO showed that MCCs displayed reduced repulsion (average time of contact +/- SD: 40.31+/-20.04 min) (Figure 6E, 6F and movie 10).

**Figure 6.**
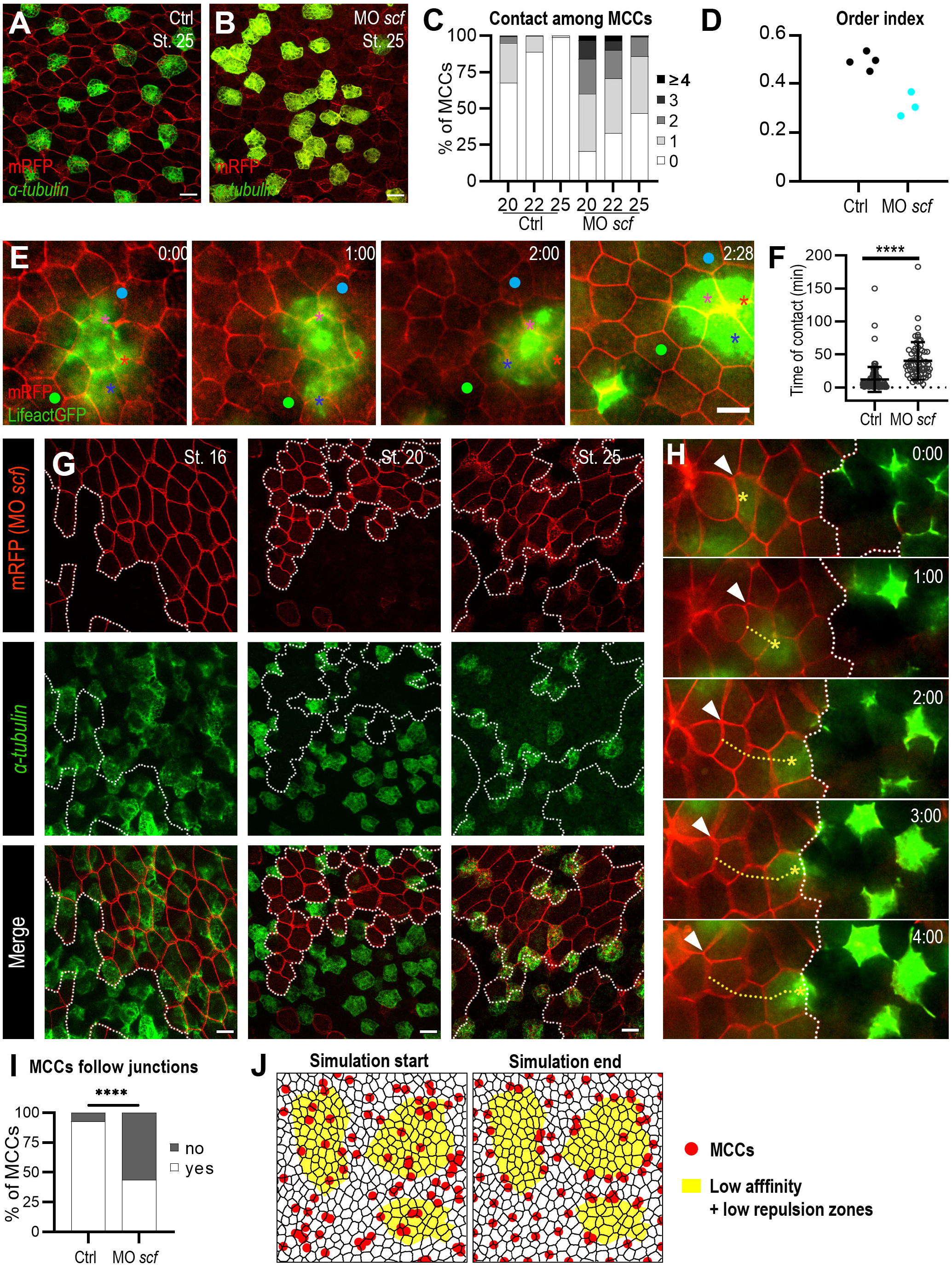
Down-regulation of the Scf signal disrupts MCC patterning. **A, B:** *α-tub* FISH (green) and RFP IF (red) revealed that compared to control embryos (**A**) MCCs were irregularly distributed in MO-ATG-scf-L/S–injected embryo (**B**) at stage 25. mRFP mRNA was used as *scf* MO tracer and to outline outer-layer cells. **C:** Quantification of MCC mutual contacts in control and *scf* MO-injected embryos. **D:** Graph showing MCC order index at stage 25 in *scf* MO-injected embryos vs control embryos. **E:** Frames from movie 10. In *scf* MO-injected embryos, LifeActGFP-positive MCCs (green) failed to properly separate from each other. mRFP (red) labelled outer-layer cell membranes. Asterisks label individual MCCs. Dots label two outer-layer cells for reference. **F:** Graph showing the duration of MCC mutual contact in control and *scf* MO-injected embryos.**** p-value < 0.0001 on a student t-test. **G:** *α-tub* FISH (green) and RFP IF (red) of embryos injected with *scf* MO at 16-cell stage and fixed at different developmental times. RFP was used as a MO tracer. MCCs were evenly distributed between MO-positive (red) and MO-negative areas at stage 16 (left column) and accumulate at the boundary between the two areas at later stages (middle and right columns). **H:** Frames from movie 11, showing a LifeActGFP-positive MCC (yellow asterisk), which left its initial position within the MO-positive area (white arrowhead) and stopped at the boundary with the MO-negative region. In contrast, MCCs outside the MO-positive area did not significantly move. The yellow dotted line shows that the tracked MCC did not follow outer-layer junctions. In all cases, scale bar is 20μm. **I:** Graph showing that a majority of MCCs did not follow outer-layer junctions in *scf* MO-injected embryos compared to control. **J:** Frames from simulation movie 12. Mathematical modelling predicted that motile MCCs tend to accumulate at boundaries between regions devoid of both repulsive and attractive signals and normal regions.

Remarkably, when *scf* MO was injected with mRFP at 16-cell stage to increase mosaicism, most MCCs at stage 25 were found at the border of, rather than within, Scf-depleted clones (Figure 6G). Analysis at stage 16 showed that MCCs were initially distributed homogeneously across *scf* MO-injected and uninjected areas, suggesting that the observed terminal distribution was due to migration of MCCs outside regions devoid of Scf. Spinning-disc video microscopy of areas straddling clonal boundaries showed that indeed, MCCs often left Scf-depleted regions and stopped migrating as soon as they encountered an uninjected outer-layer cell. Interestingly, a majority of MCCs appeared to move across *scf* MO-injected regions without following outer-layer junctions (only 43.55% of 62 analysed MCCs follow junctions) (Figure 6H, 6I and movie 11). Strikingly, *in silico* simulations revealed that MCC accumulation at clonal boundaries could only be recapitulated when both MCC mutual repulsion and affinity towards outer-layer junctions were absent from clones surrounded by normal tissue (Figure 6J and movies 12-14). This suggests that Scf might be able to convey both a repulsive and an adhesive signal.

### The Scf/Kit signalling system promotes both adhesive and repulsive functions

Our data suggest that the Scf/Kit pathway supports both mutual MCC repulsion and MCC affinity for outer-layer junctions. As a first step to test this hypothesis, we ectopically expressed the Scf-GFP chimera in MCCs by means of the α-tub promoter. *α-tub* FISH and GFP IF at stage 25 showed that this resulted in large clusters of non-injected MCCs forming around Scf-GFP-expressing MCCs (Figure 7A-C). This suggests that high-level Scf expression creates a situation where adhesion dominates repulsion. Spinning-disc video microscopy further confirmed that Scf-GFP-expressing MCCs formed closely packed groups that did not dissociate over periods of more than 3h (Figure 7D and movie 15). We conclude from these data that Scf can act as a potent adhesive cue towards Kit-expressing MCCs.

**Figure 7.**
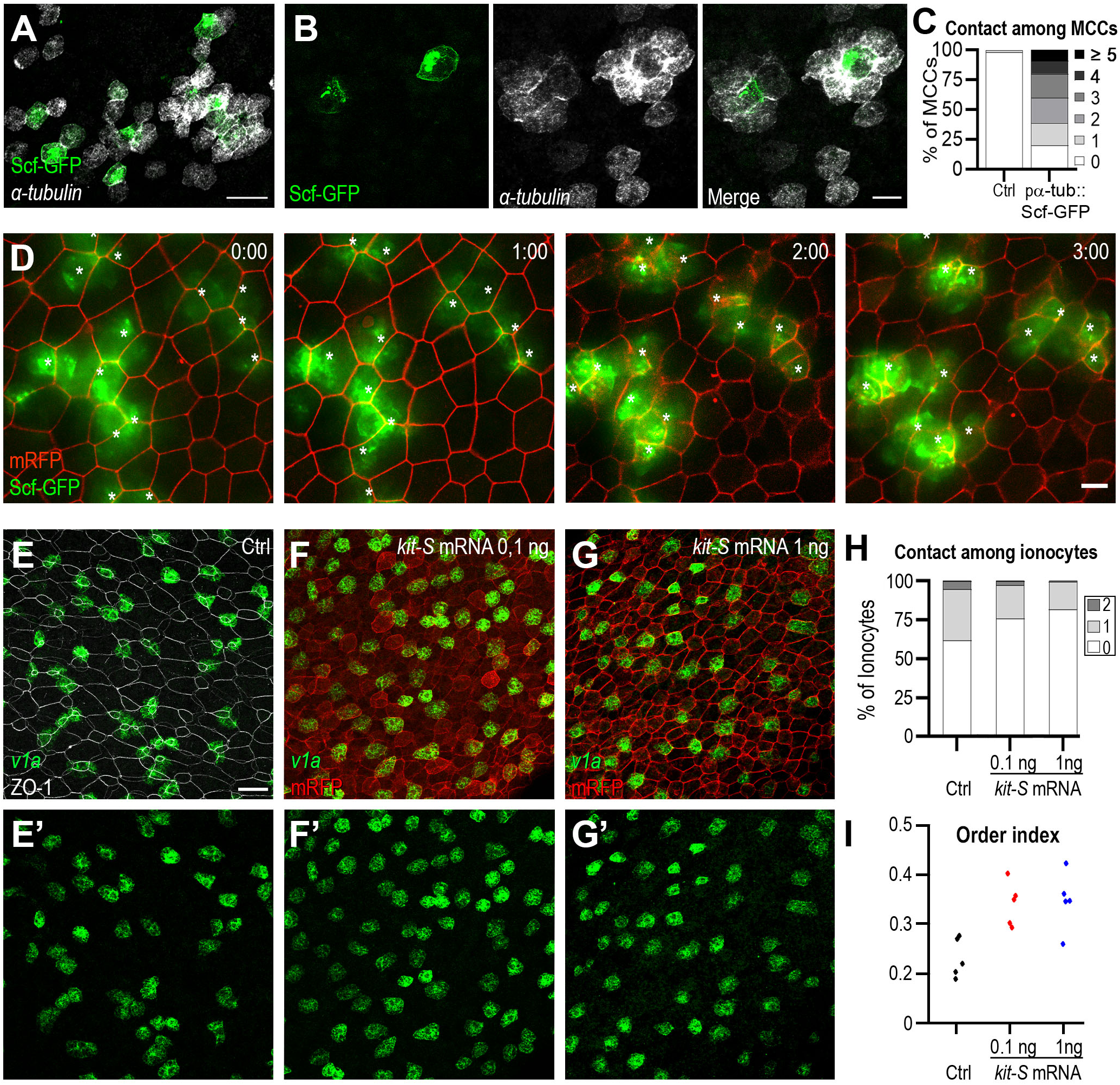
Adhesive and repulsive activities of Scf/Kit signal. **A:** *α-tub* FISH and GFP IF on stage 25 embryos injected with α-tub::Scf-GFP showed that non-injected *α-tub*-positive and Scf-GFP-positive MCCs formed aggregates. Scale bar is 50μm. **B:** A higher magnification view shows that non-injected *α-tub*-positive MCCs were tightly packed around Scf-GFP-positive MCCs. Scale bar is 20μm. **C:** Graph showing the distribution of the number of MCC mutual contacts in control and α-tub::Scf-GFP-injected embryos. **D:** Frames from movie 13 showing the stability of Scf-GFP-positive MCC clusters over time. Scale bar is 20μm. **E-G’:** Control embryos (E) and embryos injected with 0.1 or 1ng mRNA coding for full length Kit-S, fixed at stage 25 and processed for FISH with the ionocyte marker *v1a*. IF against endogenous ZO-1 (E) or against co-injected mRFP (F, G) highlighted outer-layer junctions. Kit overexpression led to more regular dispersion of ionocytes compared to control. Scale bar is 50μm. **H:** Graph showing that ionocytes established fewer mutual contacts in Kit-S-injected than in control embryos. **I:** Ionocyte order index was increased in Kit-S-injected embryos with respect to control embryos.

We then tested whether the presence of the Kit receptor was sufficient to help dispersion of non-MCC intercalating cells. To this aim, embryos were injected with an mRNA coding for full-length Kit-S and subjected to FISH with a probe against the ionocyte marker, *v1a* (Quigley et al., 2011). In control embryos, ionocytes displayed an irregular pattern of distribution and were often found in adjacent positions (Figure 7E, 7H, 7I). In contrast, *kit* RNA injection endowed ionocytes with the capacity to disperse and adopt a more regular pattern of distribution (Figure 7F-I). This last set of experiments suggest that the Ssc/Kit signalling system is sufficient to promote adhesive and repulsive functions, both necessary to establish a regular pattern of MCC distribution.

## DISCUSSION

### A cell movement-based view of *Xenopus* MCC patterning

Although *Xenopus* MCCs have been the object of extensive investigation in the past years, most studies have focused on their specification within the inner epidermal layer and their radial intercalation among outer-layer goblet cells. In contrast, much less is known on the mechanisms that control MCC distribution pattern. A pioneering study proposed that the MCC scattered pattern results from Notch-mediated lateral inhibition in the inner layer and the constraint of intercalation at outer-layer vertices (Deblandre et al., 1999). However, it was later shown that the outer layer can accommodate twice as many MCCs upon Notch inhibition, with no violation of the non-contiguity rule (Deblandre et al., 1999; Stubbs et al., 2006). This indicates that in normal condition, vertices are in large excess of incoming MCCs and that MCC spacing is not directly linked to the pattern of distribution of vertices. Our data are the first to unambiguously show that MCCs actively move within the inner layer and that motility is the main driver of their transition from a chaotic distribution, characterized by extensive inter-MCC contacts, to an orderly pattern, where MCCs are present as isolated cells, each localised in correspondence of an outer-layer vertex. In situations of severe actin or Scf/Kit pathway perturbations, we found that regular spacing between MCCs was lost despite proper intercalation. Thus, we can safely conclude that MCC pattern establishment occurs independently of radial intercalation. *In silico* simulation defined two conditions required to evenly distribute a population of internal motile cells below epithelial vertices: 1) motile cells must repel each other, thus minimizing their mutual contacts, 2) they must maximize their contacts with outer-layer junctions, so as to stabilize at vertices, which display the highest local density of junctions. Filming live MCCs in developing embryos allowed us to visualize these two behaviours and to reveal that actin cytoskeleton is involved in both. Immature MCCs in the inner layer extend unpolarized basolateral actin-rich protrusions with which they move and explore the surrounding environment. Whenever two such MCCs come into direct contact, the actin-rich protrusions involved in the contact are retracted, while others are deployed in different regions of the cell body, thus resulting in the two cells moving away from each other. Concomitantly, immature MCCs also emit apically-polarized actin-rich protrusions towards junctions between outer-layer cells. Despite the fact that both basolaterally- and apically-directed actin protrusions largely coexist within each MCC, a clear temporal shift between the two is observed: over time, the basolateral protrusive activity decreases, while the apically-directed one gains in intensity and stability (see example in Figure 2B and movie 2). This shift correlates with a progressive decrease of MCC motility within the inner layer and enhanced stabilization under individual vertices. Presumably, this transition must be asynchronous among MCCs to allow for fine tuning of the spacing pattern over time. Consistent with this idea, we found evidence that MCC specification itself may be asynchronous (Figure S1). As a result, by early tailbud stage, all MCCs end up finding a free vertex at regular intervals, to which they anchor and through which they subsequently intercalate.

### Self-organised generation of a regular cellular pattern by mutual repulsion and homing

Our data show that within the inner layer of the developing *Xenopus* epidermis, immature MCCs move in a partially random fashion, their free displacements being constrained by mutual repulsion and affinity for outer-layer intercellular junctions, and that these movements are required for the establishment of a regularly-dispersed MCC pattern. This observation prompts analogies with other instances of developmental patterns emerging from the semi-stochastic movements of cell populations. Phenomena of contact-dependent homotypic repulsion in which mutually contacting cells withdraw projections and/or change direction of movement underlie the regular distribution of Cajal-Retzius (CR) neurons (Villar-Cervino et al., 2013) and Retinal Horizontal (RH) cells (Galli-Resta et al., 2002) in mammals, pigment cells in zebrafish (Walderich et al., 2016) and haemocytes in *Drosophila* (Davis et al., 2012). The case of *Xenopus* MCC patterning shows nevertheless some specificities. Most notably, MCCs are already present throughout most of the developing epidermis at the start of the patterning process, such that their scattering relies almost exclusively on short-range “wobbling” random movements, without need for a concomitant long-range directional migration as is the case for CR neurons. As a consequence, MCCs are unpolarised and lack a well-defined leading process. Moreover, unlike *Danio* pigment cells, scattering MCCs are post-mitotic and the patterning system may not need to adjust to variations in absolute cell numbers. Finally, in sharp contrast with *Drosophila* haemocytes, which continue moving and repelling each other at every encounter, even after regular distribution is reached, dispersed MCCs become fixed by anchoring and intercalating into outer-layer vertices. Interestingly, to our knowledge, the only other described case of (semi-) orderly epithelial intercalation of a population of individually dispersed motile cells occurs during the development of the peripheral nervous system in the cephalochordate Amphioxus (Benito-Gutierrez et al., 2005; Kaltenbach et al., 2009), but the underlying mechanisms are unknown. Overall, our work reveals a new instance of self-organised tissue-wide developmental patterning, emerging from the interplay of stochastic short-range repulsive and stabilizing intercellular interactions, rather than relying on tightly controlled GRNs and classical guidance mechanisms. This is probably the most information-parsimonious strategy to optimise the distribution of a cell population whose function requires that its area of dispersion be covered completely but non-redundantly. Our identification of a molecular pathway responsible for controlling MCC dispersion, independently of cell numbers and differentiation, will make it possible to functionally explore the physiological meaning of such a pattern and the possible outcome of its perturbation.

### Implementation of *Xenopus* MCC patterning by the SCF/KIT pathway

The molecular mechanisms controlling homotypic repulsion-mediated patterning have been clearly identified in few cases only. The Eph RTKs and their ephrin ligands control the dispersion of CR neurons in the mammalian cortex (Villar-Cervino et al., 2013). Protocadherins and the immunoglobulin-superfamily proteins Dscams have been involved in the homotypic repulsion phenomena underlying mammalian RH cells tiling (Fuerst et al., 2008; Ing-Esteves et al., 2018), while the scattering of zebrafish pigment cells appears to rely on a variety of cell surface molecules, among which connexins and a potassium channel (Irion et al., 2014; Iwashita et al., 2006).

Our data show that the signal triggered by the interaction between the RTK Kit and its ligand Scf is required to establish the regularly dispersed pattern of *Xenopus* MCCs. Although this notoriously pleiotropic signalling pathway has been involved in many instances of cell migration, cell movement, cell adhesion, axonal guidance and cytoskeletal rearrangement, this is the first time that it has been shown to play a role in controlling the tissue-wide regular dispersion of a large cellular population. Remarkably, *kit* appears to be a direct transcriptional target of the multiciliogenesis master controller Multicilin/Mcidas (Kim et al., 2018; Ma et al., 2014; Stubbs et al., 2012). Such integration, through a common regulator, of fate determination and motility behaviour may explain how thousands of individual cells of the same type can self-organise to achieve a global collective pattern.

Our results are also remarkable in showing that the activity of a single receptor/ligand couple can result in the two opposite but temporally coexisting biological outcomes of MCC mutual repulsion and affinity for outer-layer junctions. Although these two phenomena might seem difficult to reconcile, the SCF/KIT signalling system has already been shown to promote apparently contrasting cell behaviours, such as migration and homing/adhesion, for example in melanoblasts (Jordan and Jackson, 2000; Wehrle-Haller and Weston, 1995), Hematopoietic Stem Cells (HSCs) (Heissig et al., 2002; Tajima et al., 1998) and Primordial Germ Cells (PGCs)(Gu et al., 2009; Gu et al., 2011; Runyan et al., 2006). Currently, the simplest and most widely accepted explanation, supported by *in vivo, ex vivo* and *in vitro* data, is that the membrane-bound and the soluble forms of the SCF ligand can both bind to and activate the KIT receptor, but with different kinetics, therefore triggering distinct, probably antagonistic signalling pathways and eliciting multiple cellular responses. In relevance to our work, soluble SCF has been shown to lead to transient activation of the KIT intrinsic TK activity, and to promote cell motility (Jordan and Jackson, 2000; Tabone-Eglinger et al., 2014; Tabone-Eglinger et al., 2012). Conversely, membrane-bound or immobilised SCF leads to formation of long-lived adhesive SCF/KIT complexes at the cell surface, independently of KIT TK activity (Tabone-Eglinger et al., 2014; Tabone-Eglinger et al., 2012). Our data show that the transmembrane Scf can be cleaved in *Xenopus* epidermis, thus raising the possibility that the dichotomy between membrane-associated and soluble Scf-dependent signals also controls the balance between adhesion and repulsion required for correct MCC patterning. It is easy to picture that the basolateral localisation of transmembrane Scf in outer-layer cells might favour/stabilise the interaction of Kit-expressing MCCs with intercellular junctions, and even more so with vertices, where the density of the Scf signal is expected to be highest. This view is supported by our observations that MO-mediated depletion of Scf or Kit altered MCC migration along outer-layer junctions. In contrast, axitinib treatment did not cause the same defect, suggesting that TK activity is dispensable for MCC/outer cell interaction, consistent with published evidence (Tabone-Eglinger et al., 2014). Our ectopic expression assay also clearly demonstrated that full-length Scf can act as a potent adhesive cue towards Kit-expressing MCCs. Incidentally, it also revealed that the patterning system cannot afford Scf to be expressed in MCCs, consistent with our double FISH data, and with transcriptomic analysis suggesting that Mcidas negatively regulates *scf* (Ma et al., 2014).

By contrast, it is less simple to imagine how soluble Scf could promote MCC mutual repulsion. One possible explanation is that soluble Scf permeating the inner layer provides a motogenic signal, as described for example for mouse melanoblasts (Jordan and Jackson, 2000), thus keeping MCCs constantly moving around and dispersing until they are anchored to a free vertex by the strongly adhesive signal of membrane-bound Scf. Alternatively, the interaction between soluble Scf and Kit might initiate a cross-talk with other molecules expressed at the MCC surface and directly responsible for the mutual repulsion response, such as ephrins and Ephs. However, the remarkable finding that the overexpression of wild-type Kit is sufficient to induce the dispersion of ionocytes suggests that the repulsive activity relies largely on the activity of Kit itself, without the need for other, MCC-specific cell surface molecules. It is thus likely that the signalling cascade connecting Kit to the cytoskeletal effectors of cell repulsion and adhesion is a rather direct and short one.

### Scf/Kit signalling and MCC actin cytoskeletal dynamics

Our data from pharmacological inhibition experiments suggest that different modes of actin polymerization, required for the formation of distinct types of cellular protrusions, control separate aspects of MCC behaviour. Following treatment with CK-666, an inhibitor of the actin-branching factor ARP2/3, MCCs present more filopodia-like extensions, show decreased mutual repulsion and major patterning defects, but no intercalation abnormalities. On the other hand, treatment with the actin bundling inhibitor SMIFH2 strongly delays intercalation without affecting patterning. The SCF/KIT pathway has been shown to control ARP2/3 activation (Mani et al., 2009) and in our model system its inhibition indeed results in defects similar to those due to CK-666 treatment: MCCs make filopodia-like protrusions and present patterning defects but are able to insert into the outer layer. Both CK-666 treatment and Scf/Kit pathway inhibition also increase the duration of MCC mutual contacts. Arp2/3 -/- fibroblasts, in which motility only relies on filopodia, show defects in mutual repulsion-dependent collective cell movements, such as those involved in wound healing (Suraneni et al., 2012).

We can thus propose a model in which the main role of the Scf/Kit pathway in *Xenopus* immature MCCs is the promotion of actin-based protrusions: Kit activation by soluble Scf would induce transient basolateral lamellipodia, required for redirecting MCC movements away from each other after mutual contact; membrane bound SCF would stabilize the more persistent apical protrusions required for anchoring MCCs to the vertices. Such a model warrants future validation by cell biological and biochemical approaches.

## Conclusions

We show here that, during development, seemingly complex and highly ordered cell patterns can be obtained by the combination of two relatively simple cell behaviours, without need for a highly regulated patterning system. Moreover, deployment of the same signalling system, presumably activated with different kinetics by soluble and membrane-anchored forms of a single ligand, can control both behaviours. This further stresses the importance of self-organising systems in developmental biology.

## ACKNOWLEDGMENTS

AC was supported by a doctoral fellowship from the French Ministry for Higher Education, Research and Innovation (MESRI). CR was supported by a postdoctoral fellowship from the Turing Centre for Living Systems (CENTURI). The project leading to this publication has received funding from Excellence Initiative of Aix-Marseille University - A*MIDEX, a French “Investments for the Future” program (Projet ‘Pépinière d’Excellence 2017 to AP). The work was also supported by funding from Cancéropôle PACA (Programme Emergence 2017to AP), the Fondation pour la Recherche Médicale (DEQ20141231765, EQU201903007834 to LK), and the Association pour la Recherche contre le Cancer (PJA 20141201815 to LK). Imaging was performed on PiCSL-FBI core facility supported by the French National Research Agency through the “Investments for the Future” program (France-BioImaging, ANR-10-INBS-04). We are indebted to E. Bazellières, J. Christian, R. Harland, C. Kintner, A. Miller, M. Ogawa, O. Rosnet and J-P. Tassan for the gift of plasmids. We thank F. Roguet for *Xenopus* husbandry. We warmly thank Anna Adamiok, Marie Cibois and Marie Zilliox for preliminary experiments, Marc Billaud, Claudio Collinet and Laurence Röder for insights on the project, past and present members of the Kodjabachian laboratory, in particular Camille Boutin and Olivier Rosnet, for discussions, help and technical support, Elsa Bazellières and Vincent Bertrand for insightful comments on the manuscript.

## AUTHOR CONTRIBUTIONS

Conceptualisation: AC, LK, AP. Methodology: AC, RC, CR, VT, AP. Software: RC, CR. Formal analysis: AC, RC, CR. Investigation: AC, RC, CR, VT, AP. Validation: AC, CR, VT, RC, LK, AP. Resources: LK. Supervision: LK, AP, RC. Project administration: LK, AP. Funding acquisition: LK, AP, RC. Writing - original draft: AC, AP. Writing - review and editing: LK, AP.

## DECLARATION OF INTERESTS

The authors declare no competing interests.

## STAR METHODS

### Ethics statement

All procedures were performed following the Directive 2010/63/EU of the European parliament and of the council of 22 September 2010 on the protection of animals used for scientific purposes. Experiments on *X. laevis* were approved by the ‘Direction Départementale de la Protection des Populations, Pôle Alimentation, Santé Animale, Environnement, des Bouches du Rhône’ (agreement number F 13 055 21).

### Embryo culture and injection

Ovulation was stimulated in *X. laevis* adult females from NASCO (https://www.enasco.com) by injection of (800 units/animal) Human Chorionic Gonadotropin (Chorulon^R^). On the following day, eggs were recovered by squeezing, fertilized *in vitro* with sperm from NASCO males, de-jellied in 2% cysteine hydrochloride (pH 8.0) and washed, first in water, then in 0.1X MBS (Modified Barth’s Saline). Embryos were kept in 0.1X MBS at 13°C, 18°C or 23°C until they reached the stage suitable for injection (8-cell or 16-cell), then transferred in 4% Ficoll in 1X MBS. Injections were performed using needles made from pulled glass capillaries and mounted on a Nanoject II injector (Drummond). Embryos were injected in animal ventral blastomeres to target the non-neural ectoderm. One blastomere (in 8-cell stage embryos) or two contralateral blastomeres (in 16-cell stage embryos) were injected, according to the desired degree of mosaicism.

### Plasmids

| Plamid name | backbone | linearized<br>with | amount<br>injected/blastomere | Reference |
| --- | --- | --- | --- | --- |
| pαTub::GFP | pCS2+ | NotI | 50-60pg | Stubbs et al. |
|  |  |  |  | 2006 |
| pαTub::RFP | pCS2+ | NotI | 50-60pg | Made in the lab |
| pαTub::LifeActGFP | pCS2+ | NotI | 50-60pg | |
| pαTub::LifeActRFP | pCS2+ | NotI | 50-60pg | |
| pαTub::EGFPp41 | pCS2+ | NotI | 30-50pg | |
| pαTub::Scf-GFP | pCS2+ | NotI | 50-60pg | |
| pCS107-Scf-L-GFP | pCS107 | NotI | // |  |
| pαTub::MOresKit-S | pCS2+ | NotI | // | |
| pCMV-MOresKit-S | pCMVSport | NotI | // |  |

All αTubulin promoter-based constructs were generated from pαTub::GFP (a gift from C. Kintner), either with the In-Fusion^R^ HD Cloning Kit (Takara Bio) or by standard restriction enzyme-mediated cloning.

For In-Fusion cloning, the pαTub:: backbone, excluding the GFP ORF, was PCR amplified using the Phusion^R^ High-Fidelity DNA Polymerase (New England BioLabs, #M0530) and the primers pαtubfor:5’-TCTAGAACTATAGTGAGTCGT-3’ and pαtubrev:5’-CGACCGGTGGATCTGTTGTTGG-3’.

To generate pαTub::RFP, the RFP ORF was PCR amplified from the pCS2+mRFP plasmid (a gift from O. Rosnet) with the Phusion^R^ High-Fidelity DNA Polymerase and the primers InFuRFPfor:5’-CAGATCCACCGGTCGCATGGCCTCCTCCGAGGACGT-3’ and InFuRFPrev:5’-CACTATAGTTCTAGATTAGGCGCCGGTGGAGTGGCG.

To generate pαTub::LifeActGFP, the LifeActGFP ORF was PCR amplified from the pmEGFP-Lifeact-7 plasmid (a gift from E. Bazellières) with the Phusion^R^ High-Fidelity DNA Polymerase and the primers InFuLifeactfor:5’-CAGATCCACCGGTCGTATGGGTGTCGCAGATTTGAT-3’ and InFuLifeactGFPrev:5’-CACTATAGTTCTAGATTACTTGTACAGCTCGTCCA-3’.

To generate pαTub::LifeActRFP, the LifeActRFP ORF was PCR amplified from the pmRuby-Lifeact-7 plasmid (a gift from E. Bazellières) with the Phusion^R^ High-Fidelity DNA Polymerase and the primers InFuLifeactfor:5’-CAGATCCACCGGTCGTATGGGTGTCGCAGATTTGAT-3’ and InFuRFPrev:5’-CACTATAGTTCTAGATTAGGCGCCGGTGGAGTGGCG.

To generate pαTub::EGFPp41, the EGFPp41 ORF was PCR amplified from the EGFP-p41-Arc plasmid (a gift from M. Ogawa) with the Phusion^R^ High-Fidelity DNA Polymerase and the primers InFuEGFPfor:5’-CAGATCCACCGGTCGTATGGTGAGCAAGGGCGAGGAG-3’ and InFup41rev:5’-CACTATAGTTCTAGATCACTTAATCTTCAGATCTTT-3’.

In all the above cases, the vector and insert PCR products were run on agarose/TAE gel, purified with the Macherey-Nagel™ NucleoSpin™ Gel and PCR Clean-up kit, then recombined using the In-Fusion^R^ HD Cloning Kit according to the manufacturer’s instructions.

To make the pCS107-Scf-L-GFP construct, the pCS107-XlSteel1 plasmid (a gift of R. Harland), containing the Xl-Scf-L cDNA, was PCR amplified using the Phusion^R^ High-Fidelity DNA Polymerase and the primers Steel1fus-for:5’-GCCACAAATCCCCCTTGTAAAGT-3’ and Scf1-3’endfus-rev:5’-TATAACACCGACATCAGTTGTGG-3’, in order to generate a linearized backbone lacking the Scf-L stop codon and 3’UTR. The GFP ORF was PCR amplified from the pαTub::GFP construct using the Phusion^R^ High-Fidelity DNA Polymerase and the primers Scf1-GFPfus-for:5’-GATGTCGGTGTTATAGTGAGCAAGGGCGAGGAGC-3’ and GFP3’Scf1fus-rev:AGGGGGATTTGTGGCTTACTTGTACAGCTCGTCACT-3’. Vector and insert PCR products were run on agarose/TAE gel, purified with the Macherey-Nagel™ NucleoSpin™ Gel and PCR Clean-up kit, then recombined using the In-Fusion^R^ HD Cloning Kit according to the manufacturer’s instructions.

The pCS107-Scf-L-GFP plasmid was used both as template for making Scf-GFP mRNA and to generate the pαTub::Scf-GFP construct. To this aim, the Scf-L-GFP ORF was PCR amplified from pCS107-Scf-L-GFP using the Phusion^R^ High-Fidelity DNA Polymerase and the primers αtubSCFGFPfor:5-CAGATCCACCGGTCGTATGAAGAAGACAAAAACTTGG-3’ and InFuLifeactGFPrev:5’-CACTATAGTTCTAGATTACTTGTACAGCTCGTCCA-3’. The gel-purified PCR product was recombined into the pαTub:: backbone using the In-Fusion^R^ HD Cloning Kit.

To generate the pCMV-MOresKit-S plasmid, the *kit-S* ORF was PCR amplified from the Dharmacon/GE clone 4030854 (*Xenopus laevis kit-b*), using the Phusion^R^ High-Fidelity DNA Polymerase and the primers αtubMOresKitSfor:5-CAGATCCACCGGTCGTGCTACA**ATG**AG**<u>T</u>**CA**<u>C</u>**AT**<u>C</u>**CC**<u>T</u>**TT**<u>T</u>**CT**<u>G</u>**AC**<u>T</u>**GTCAGG -3’ (in bold, the Kit-S start codon, in bold underlined the seven silent mutations introduced to generate mismatches with MO-ATG-kit-S) and αtuBKitSrev:5’-CACTATAGTTCTAGATCAGCCGTCGCTGTTCATCAACAG-3’. The gel-purified PCR product was recombined into the pαTub:: backbone using the In-Fusion^R^ HD Cloning Kit, thus creating the pαTub::MOresKit-S plasmid. This was subsequently cut with AgeI and NotI and the MOresKit-S ORF-containing fragment was ligated into AgeI/NotI cut pCMVSport6.

All constructs were made using the StellarTM Competent Cells (Clontech) transformed and grown according to the manufacturer’s instructions. Small- and medium-scale plasmid preparations were performed with Macherey-Nagel™ NucleoSpin™ Plasmid QuickPure™ and NucleoBond™ Extra Midi kits.

### mRNA synthesis

Sense mRNAs were synthesized from linearized plasmids with the Ambion mMessage mMachine kit® (Life Technologies) according to the details provided in the table below, then purified with Macherey-Nagel™ NucleoSpin™ RNA Clean-up kit. After determination of the concentration, aliquots were kept at −80°C.

| Plasmid name | Vector | Linearised<br>with | Transcribed<br>with | Amount<br>injected<br>/blastomere | Reference |
| --- | --- | --- | --- | --- | --- |
| pCS2+mRFP | pCS2+ | NotI | Sp6 RNA pol | 0,5 ng |  |
| pCS2+GFPgpi | pCS2+ | NotI | Sp6 RNA pol | 0,5ng | Chartrain et<br>al., 2003 |
| pCS107-Scf-L-GFP | pCS107 | NotI | Sp6 RNA pol | 1 ng | Made in the lab |
| pCS2+RFP-ZO1 | pCS2+ | NotI | Sp6 RNA pol | 1ng | Higashi et al., 2016 |
| pCS2+dnKit | pCS2+ | NotI | Sp6 RNA pol | 0,5ng | Goldman et al., 2006 |
| pCMV-dnKit-GFP | pCMVSport6 | NotI | Sp6 RNA pol | 0,5g | Made in the lab |
| pCMV-MOresKit-S | pCMVSport6 | NotI | Sp6 RNA pol | 0.1-1ng | Made in the lab |

### Morpholinos

Morpholinos (MOs) were purchased from GENE TOOLS®. ATG-kit-L-MO and ATG-kit-S-MO were injected together at 10ng/MO/blastomere. kit-S-e2i2spl-MO and kit-L-e2i2spl-MO were injected together at 15ng/MO/blastomere. ATG-scf-L/S-MO was injected at 10ng/blastomere and scf-L-e2i2spl-MO at 20ng/blastomere. The blue-tagged version of ATG-scf-L/S-MO was injected at 20ng/blastomere.

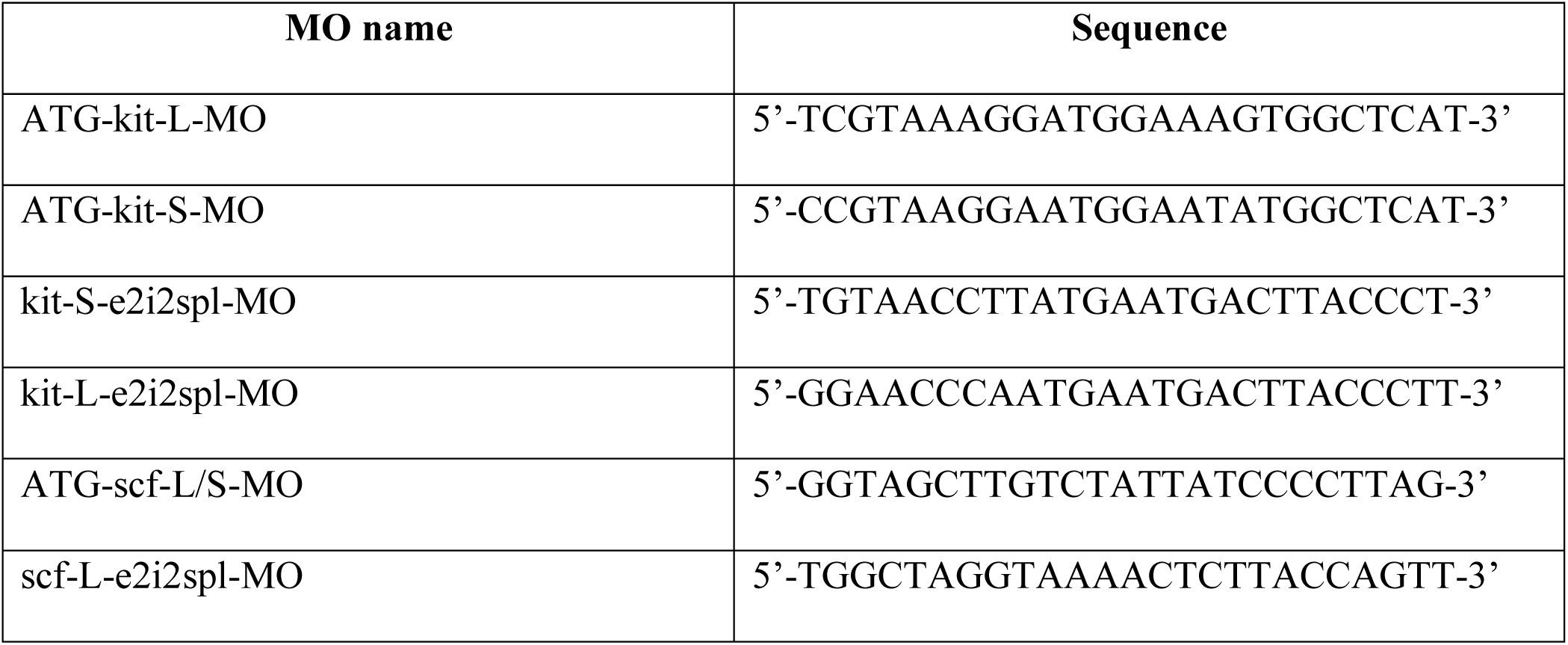

### Pharmacological inhibitors

Stock solutions of pharmacological inhibitors (all from Sigma-Aldrich®) were prepared in DMSO and added to embryos cultured in MBS 0,1X at the stages and concentrations and for the times shown in the following table. Cycloheximide treatment was perform by incubating embryos at stage 14 in MBS 0,1X containing 300µM cycloheximide two hours before live imaging.

| Drugs | Source | Reference | Concentration<br>in 0,1X MBS | Incubation time and<br>stage |
| --- | --- | --- | --- | --- |
| Axitinib | Sigma-Aldrich | AG-013736 | 50µM | From stage 12 |
| CK-666 |  | SML0006 | 100µM |  |
| SMIFH2 |  | S4826 | 50µM |  |
| Cycloheximide |  | C7698 | 300µM | From stage 14 |

### Apoptosis detection

The CellEvent™ Caspase-3/7 Green Detection Reagent (Invitrogen™) was dissolved in DMSO and added to stage 14 control or cycloheximide-treated embryos in MBS 0,1x at 20µM final concentration.

### *In situ* hybridization and immunostaining

Whole-mount chromogenic In Situ Hybridization (ISH) and whole-mount Fluorescent In Situ Hybridization (FISH) were performed as previously detailed (Castillo-Briceno and Kodjabachian, 2014; Marchal et al., 2009). Embryos were fixed in MEMFA (0.1 M MOPS pH 7.4, 2 mM MgSO4, 1 mM EGTA, 3.7% v/v formaldehyde) 1h30 at room temperature or overnight at 4 °C then dehydrated o/n in 100% methanol at −20 °C. For FISH on section, embryos were fixed in MEMFA, stored in methanol o/n at −20 °C, rehydrated in PBT (PBS + Tween 0.1% v/v), treated with triethanolamine 100mM and acetic anhydride, incubated in increasing sucrose concentrations, embedded in OCT (VWR Chemicals), cut in 12 µm-thick cryosections and stored at −80°C. Before hybridization, embryos were rehydrated in PBT (PBS+0.1% Tween20), treated with Proteinase K (8 min at 2 µg.ml^−1^), then, for FISH, the endogenous peroxydase activity was blocked by incubation with H_2_O_2_ 3% in PBS for 20 min. For single staining, RNA probes were labeled with digoxigenin-dUTP (Roche). For double staining, various combinations of digoxigenin/fluorescein-labeled riboprobes were used. For each hybridization, the following amounts of riboprobes were used: *α-tubulin* 1ng; *v1a* 40 ng; *foxj1* 100ng; *kit* 200ng; *scf-L* 100ng to 300ng for embryos at stage 11-16 and stage19-20, respectively. Hybridization was performed overnight at 60 °C. After hybridization, the embryos were washed at increasing stringency in SSC/0.1% CHAPS, rinsed extensively in MABX (Maleic Acid Buffered solution +0.1% Triton X), then the digoxigenin-labelled probe was revealed through incubation with a sheep anti-DIG antibody conjugated to HRP (POD) (Roche, 1:500). On the third day, embryos were extensively washed in MABX, then staining was revealed using Tyramide Signal Amplification—TSA TM Plus Cyanine 3/5/Fluorescein System (PerkinElmer®). This reaction was then blocked in a bath of 2% H2O2 for 20 min.

Following FISH labeling, immunostaining was performed by incubating the embryos in MABX-BR2%, 15% FBS with the antibodies listed in the following table. After extensive washes in MABX, embryos were flattened by cutting along the antero-posterior axis and mounted in Mowiol (Sigma-Aldrich) for confocal imaging.

| Antibody | Species | Source | Dilution |
| --- | --- | --- | --- |
| ZO1-TJP1 | Mouse IgG1 | Thermo<br>Scientific | 1:200 |
| GFP | Chicken | Aves Labs | 1:500 |
| RFP | Rat monoclonal<br>IgG2a | Chromotek | 1:500 |
| Alexa Fluor 568 anti-rat | Goat | Molecular | 1:500 |
| Alexa Fluor 488 anti-chicken | Goat | Probes | 1:500 |
| Alexa Fluor 647 anti-mouse<br>IgG1 | goat |  | 1:800 |

### Confocal and electron microscopy

To limit the movement of the embryo during live imaging, control or treated stage 14-15 embryos were embedded in 0,8% LMP agar in 0,1X MBS within a small chamber made with two coverslips and silicon grease. Time-lapse imaging was done at 23°C starting at stage 14-15 and for a duration of 2 to 8 hours on a Nikon Roper spinning disc Eclipse Ti inverted microscope using a 20X_objective (for apoptosis recording), or 40X/60X _1.25 N.A water-immersion objective (for all the other experiments). Image acquisition was performed with the Meta-Morph software. 20µm-deep Z sections were acquired every 0,5 to 2min in 0,5 to 0,7µm steps, averaging 2. To compensate for the fluorescence intensity variations across different injected embryos, the laser power was adjusted for each experiment. Excitation wavelengths were 488nm, 561nm and 445nm for GFP- and RFP-tagged protein and for blue-tagged MO, respectively.

Fixed whole embryos and sections were examined on Zeiss LSM 510 and 880 confocal microscopes. Four-color confocal Z-series images were acquired using sequential laser excitation, converted into single plane projection and analysed using ImageJ/FIJI software (see image analysis section).

For Scanning Electron Microscopy (SEM), stage 27 control or treated embryos were fixed for 4h in 3% glutaraldehyde in 0.1 M phosphate buffer pH 7.4 (19 mL monosodium phosphate 0.2 M and 81 mL disodium phosphate 0.2 M), washed in phosphate buffer and filtered bi-distilled water, progressively dehydrated in ethanol at 25%, 50%, and 70% for 30 min each time, then stored in fresh ethanol 70% at 4 °C o/n. Embryos were further dehydrated with vigorous agitation once in 90% ethanol, and twice in 100% ethanol, for 30 min each, then subjected to CO_2_ critical point desiccation (CPD030, Balzers) at 40 °C and 75-80 bars. Finally, samples were sputter-coated with gold (vacuum 1 × 10–12 Torr, beam energy 3–4 keV) and stuck on the support with conductive glue for immediate SEM digital imaging (FEI TENEO) of the skin epidermis.

### Image analysis

All images were processed with the imageJ/Fiji free software.

Z-stacks from confocal video microscopy were converted into single plane projection by maximum intensity or sum intensity projection. Each channel was processed separately with the optimized z-section corresponding to the outer/inner layer cell staining. All visible MCCs were manually tracked with the ‘manual tracking’ plugin. Cell contact events were estimated by counting the time between the first and the last non-ambiguous contact time points. Cell contacts already established at the beginning of filming or persisting beyond its end were excluded from quantification. To characterize the migratory path of MCCs relative to outer-layer cell junctions the following criteria were applied: MCC emits protrusions in correspondence of outer-layer cells junctions; overlap between MCC shape and outer-layer cells junctions; correlation between direction of MCC migration and orientation of the outer-layer cells junctions. Cells already engaged in radial intercalation with permanent contact with vertices and ambiguous cases were removed from quantification.

Analysis of the apical-basal global cell-shape was done through maximum intensity projections of resliced z-stacks following X,Z and Y,Z orientations with 0,5µm interval from images at the indicated time point. Analysis of Scf-GFP/ZO-1RFP signal localization was done on resliced z-stacks from z-series at 0.2µm intervals, taken along chosen intercellular junctions. 3D-projection of intercalating MCCs probing junctions and vertices was made using the ‘clear volume’ plugin. To check for the depletion of Scf-GFP protein by a blue-fluorescent tagged version of ATG-scf-L/S-MO in live we proceeded as follows: mRNAs coding for Scf-GFP and mRFP were co-injected in the two animal ventral blastomeres at stage 8-cell stage, then blue-tagged ATG-scf-L/S-MO was injected in two of the four animal ventral blastomeres at 16-cell stage. GFP, RFP and blue fluorescence were recorded at stage 16.

Images from fixed whole embryos and sections obtained with 510 and 880 LSM confocal microscopes were converted into single plane projection by maximum intensity projections. To calculate MCC and ionocyte mutual contacts and order indexes, as well as the relation of MCCs to vertices and their intercalation, the outlines of individual *α-tubulin*- or *v1a*-positive cells were manually drawn from maximum intensity projections of 20X 450×450µm images using Wacom® Intuos Pro graphic tablet and FIJI ‘ROI manager’ tool. Mutual contacts and overlapping to ZO-1-positive vertices were manually defined. MCCs were considered intercalated when their apical surface was clearly outlined by a ZO-1 signal. Ambiguous cases were classified as ‘non defined’ (nd). To calculate the Order Index, cell areas and centroids were defined and the data processed as described in the section ‘Mathematical Modelling’.

### Western blots

Animal caps were obtained by manual dissection from stage 10 embryo in 1X MBS (Sive et al, 2000) and kept in 0,5X MBS until matched control embryos reached stage 17-18, then snap-frozen in liquid nitrogen and stored at −80°C or immediately lysed by boiling for 5min in 50mM Tris-HCl (pH 7.5), 2% w/v SDS, 1mM DTT. After addition of 5x Bromophenol blue/Glycerol, samples were loaded on Acrylamide PAGE gels and run using the Hoefer™ Mighty Small™ II Mini system with Tris/Glycine/SDS buffer. Transfer to nitrocellulose membranes was performed in Tris/Glycine/Ethanol buffer using the Hoefer™ TE22 Mini Tank Blotting Unit. Membranes were rinsed with water, stained with Ponceau Red, washed three times for 10min each time in TBS-T (20mM Tris-HCl, 150mM NaCl, pH 7.5, 0.05% v/v), blocked for 1h in TBS-T+5% (w/v) non-fat dry milk, then incubated with a rabbit anti-GFP antibody (Torrey Pines Biolabs, TP401, 1:1000 in TBS-T + non-fat dry milk) overnight at 4°C. After 4 washes (15min each) in TBS-T, membranes were incubated for 1h at room temperature with a goat anti-rabbit-HRP-conjugated antibody (Invitrogen, 1:5000 in TBS-T + non-fat dry milk) then washed four times (15min each) in TBS-T. The signal obtained from enhanced chemiluminescence (Western lightning ECL Pro, Perkin Elmer) was detected with MyECL Imager (Life Technologies).

### PCR

Whole embryos or animal caps were snap frozen at different stages and stored at −80 °C. Total RNAs were purified with the Qiagen RNeasy kit (Qiagen). RT reactions were carried out using iScript™ Reverse Transcription Supermix (BIO-RAD). PCRs were carried out with GoTaq® G2 Flexi DNA Polymerase (Promega). Amplification of exon6 was performed using the primers scf-L-ex6for: 5’-GGACCTTGTACCATGCCTGC-3 and scf-L-ex6rev: 5’-ATTCCTCTGCCAGGTCTGGA-3 for *scf-L*; scf-S-ex6for: 5’-TCGGCCTCTTCTTTGTATCG-3’ and scf-S-ex6rev: 5’-GTACAGTACAGTGTTAATAG-3’ for *scf-S*. To check for the efficiency of kit-S-e2i2spl-MO, the following primers were used: check-splMOkitS1for: 5’-ACCATGAACTGGACTTACCTGTGG-3’ and check-splMOkitLSrev: 5’-CTTCAATGTCCCACTCTGATTTCC-3’. To check for the efficiency of kit-L-e2i2spl-MO the following primers were used: check-splMOkitL1for: 5’-TCCTTTACGATCACAATGAGCTGG-3’ and check-splMOkitL1rev: 5’-TTACATGAATAGAGAATGTGCTGC-3’.

### Statistical analysis

Graphs were done with Graphpad Prism8 software as well as statistical analysis when indicated using student t-test. Time of contact among MCCs was compared using unpaired parametric student t-test and junction following behaviour by Fisher test. The number of samples analysed in all graphs is presented in Table S1.

### Mathematical modelling

#### Geometry

The outer layer, considered as a static canvas that can influence the behaviour of two-dimensional *N*_*cells*_ MCCs (red disks), is extracted from the segmented image of a ZO-1 IF of a stage-14 control embryo. The system dynamics were modelled using the effective energies associated with the two mechanisms of MCC mutual repulsion and affinity for outer layer intercellular junctions. The evolution from a random initial state to the final state was guided by an algorithm imposing a progressive decrease of the system total energy. Mathematical expressions for the repulsion and affinity energies were chosen in correspondence with the qualitative behaviours observed in experiments.

#### Energies

To account for mutual repulsion, a pair-repulsion energy with an exponential decay on a typical length scale equal to the MCC radius *r*_*cell*_ was chosen. To prevent the overlap among neighbouring MCCs, the repulsion energy was supplemented with a hard-core corresponding to the minimal allowed distance between two cells. Based on experimental observations showing that MCC overlap does not exceed half their size, the hard core was defined as *r*_*cell*_. To ensure that two nearby MCCs undergo a local energy gradient and to prevent aberrant configurations where two MCCs remain stuck with a distance smaller than *r*_*cell*_, the hard-core energy decreases between *r*_*ij*_ = 0 and *r*_*cell*_.

Thus, the pair-repulsion energy reads 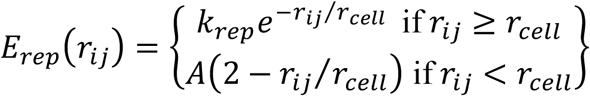, where *r*_*ij*_ is the distance between the centers of MCCs *i* and *j, r*_*cell*_ is the MCC radius and *k*_*rep*_ the repulsion constant (positive). The constant *A* of the hard-core energy is chosen large (*A* = 100), so that the probability for a MCC to be at a distance lower than *r*_*cell*_ from another MCC is very small (*e*^−100^ ≃ 10^−44^).

Affinity for junctions was accounted for with a negative energy term proportional to the overlap between MCCs and outer-layer junctions, calculated as the total pixel intensity of the junction network covered by MCCs. The affinity energy of a cell *i* is then 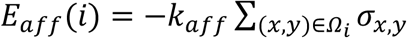 where *k*_*aff*_ is the affinity constant (positive), *Ω*_*i*_ is the set of pixel positions covered by the MCC *i* and *σ*_*x, y*_ is the value of the pixel at the position (*x, y*) of the epithelial canvas. The minus sign ensures that the energy decreases when the total pixel intensity covered by the MCC *i* increases.

To build the epithelial canvas used to calculate the affinity energy, a Gaussian blur of a size equal to a MCC diameter was applied to the segmented epithelium matrix. This considerably fastens the simulations by directing MCCs towards junctions thanks to an energy gradient, without changing the final state. Note that this could also reflect that regions lining junctions are mechanically more favourable, as the height of the epithelial cells is minimal at junctions.

#### Dynamics

The total energy of the system is the sum of the repulsion and affinity energies of all the MCCs *E*_*tot*_ = Σ_*i*_ *E*_*aff*_ (*i*) + Σ_*i*≠*j*_ *E*_*rep*_ (*r*_*ij*_) where *i* and *j* refer to MCCs. Monte-Carlo simulations of the system were performed with a Metropolis algorithm to impose the energy minimization constraint. At each time step of the simulation a random displacement was computed and attributed to each MCC. This displacement corresponds to a diffusive motion with a diffusion constant yielding typical jumps of one-tenth of a cell-radius distance. Moves were accepted or rejected following a Metropolis algorithm depending on the value of Δ*E*_*tot*_ = *E*_*tot*_(*after*) − *E*_*tot*_(*before*), the difference between the system total energy after and before each move. If the total energy has decreased (Δ*E*_*tot*_ < 0) the move is accepted. If the total energy has increased (Δ*E*_*tot*_ > 0) the move is rejected in most cases but has a nonzero probability to be accepted if the increase is of the order of the thermal energy *k*_*B*_*T* (*k*_*B*_ is the Boltzmann constant and *T* the temperature, both set to 1 in our simulations), with a probability given by exp(−Δ *E*_*tot*_ /(*k*_*B*_*T*)). The virtual-time of the simulation is increased by one after *N*_*cells*_ such dynamical steps (one Monte-Carlo time step). The simulation was run for a Monte-Carlo time sufficient for the system to reach a stationary final state. Depending on the respective weight of the energy constants *k*_*rep*_ and *k*_*aff*_ final states with different qualitative patterning of the MCCs were generated. Let alpha be the ratio between energy constants, α = *k* _*rep*_ /*k*_*aff*_. When α ≿ 10^7^, the total energy of the system *E* _*tot*_ is dominated by the repulsion energy. When α ≾ 10^3^, the total energy of the system is dominated by the affinity energy.

#### Quantification

In order to have a more quantitative description of the system state, two indices were defined, to quantify the MCC spatial order and their localization at junctions, respectively.

The order index *I*_0_ was computed from Delaunay tessellation of MCC positions. The position of each MCC is computed as a single point, located at its centroid. Systems with different MCC densities *ρ*_*cells*_ (defined as the ratio between the total area occupied by MCCs and the available area of the epithelium) were distinguished, since the number of possible final MCC organisations increases with the available space. The order index *I*_0_ is a measure of the narrowness of the distribution of areas *A*_*Del*_ of Delaunay triangles while accounting for the available space 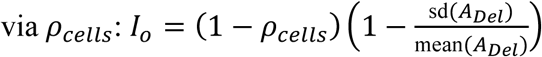,

*I*_0_ is minimal when the standard deviation of Delaunay areas is of the order of its mean and is large when the standard deviation is small compared to the mean. Therefore *I*_0_ is maximal when MCCs are perfectly ordered and minimal when they are most disordered. Note that this definition of order was used both for simulations and experimental data.

The covering index *I*_*c*_, representing the extent of overlapping between outer-layer junctions and MCCs, was computed as the total pixel intensity of the epithelium covered by MCCs, 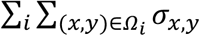, minus a noise term. This noise term, corresponding to the background value of covered pixels in a purely random configuration of MCCs, was computed by averaging the pixels *σ*_*x,y*_ of the whole epithelium and multiplying this value by the cell surface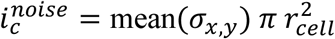. The covering index was normalized by a term accounting for the maximal possible extent of overlapping of junctions by one MCC. This was obtained by finding the maximal possible overlapping value for a cell *C* scanning among all possible positions (*x*_*C*_, *y*_*C*_) of its center 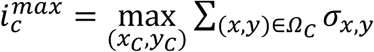.

Thus the covering index is 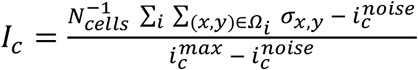.

The index is thus minimal when the overlap of MCCs with junctions does not exceed the overlap obtained from a random scattering of MCCs. It is maximal when the overlap is the highest possible.

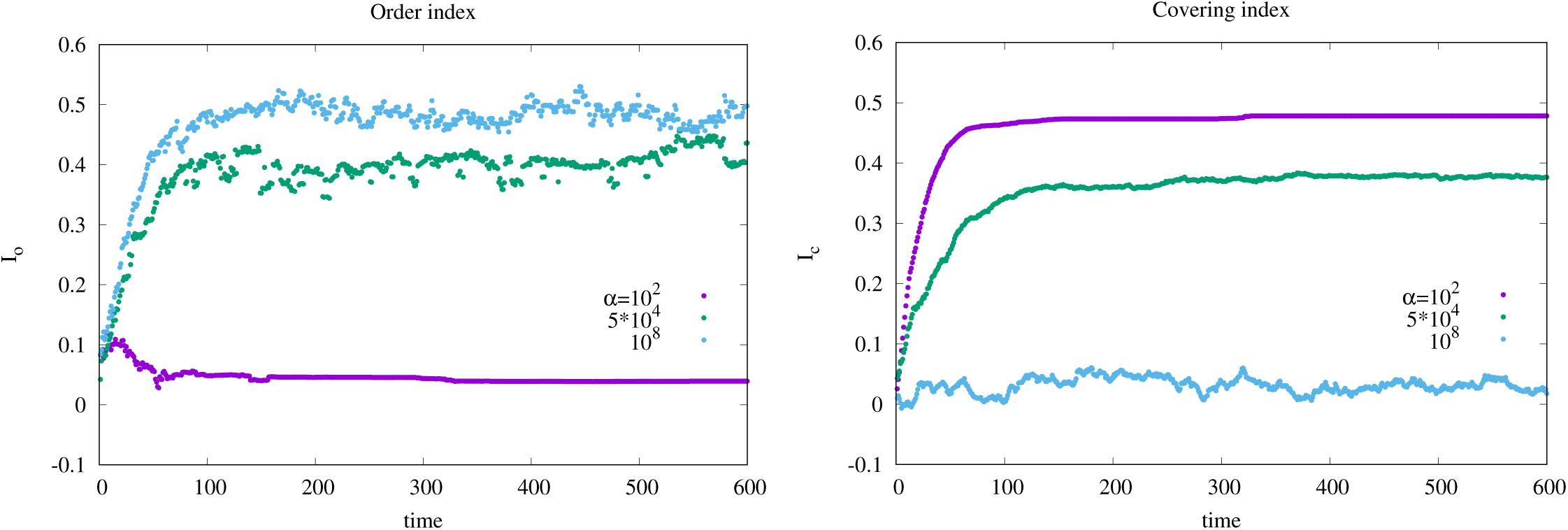

### Order and covering index evolution with time for different values of α = *k*_*rep*_/*k*_*aff*_

In the repulsion dominated regime (α=10^8^) MCCs evolve toward a strict ordering regardless of junctions hence the order index is large while the affinity index is small. In the affinity-dominated regime (α=10^2^), MCCs finally localize under junctions/vertices regardless of other MCCs, yielding a large affinity index while the order index remains small. In the intermediate regime (α=5×10^4^) the two indices take intermediate values.

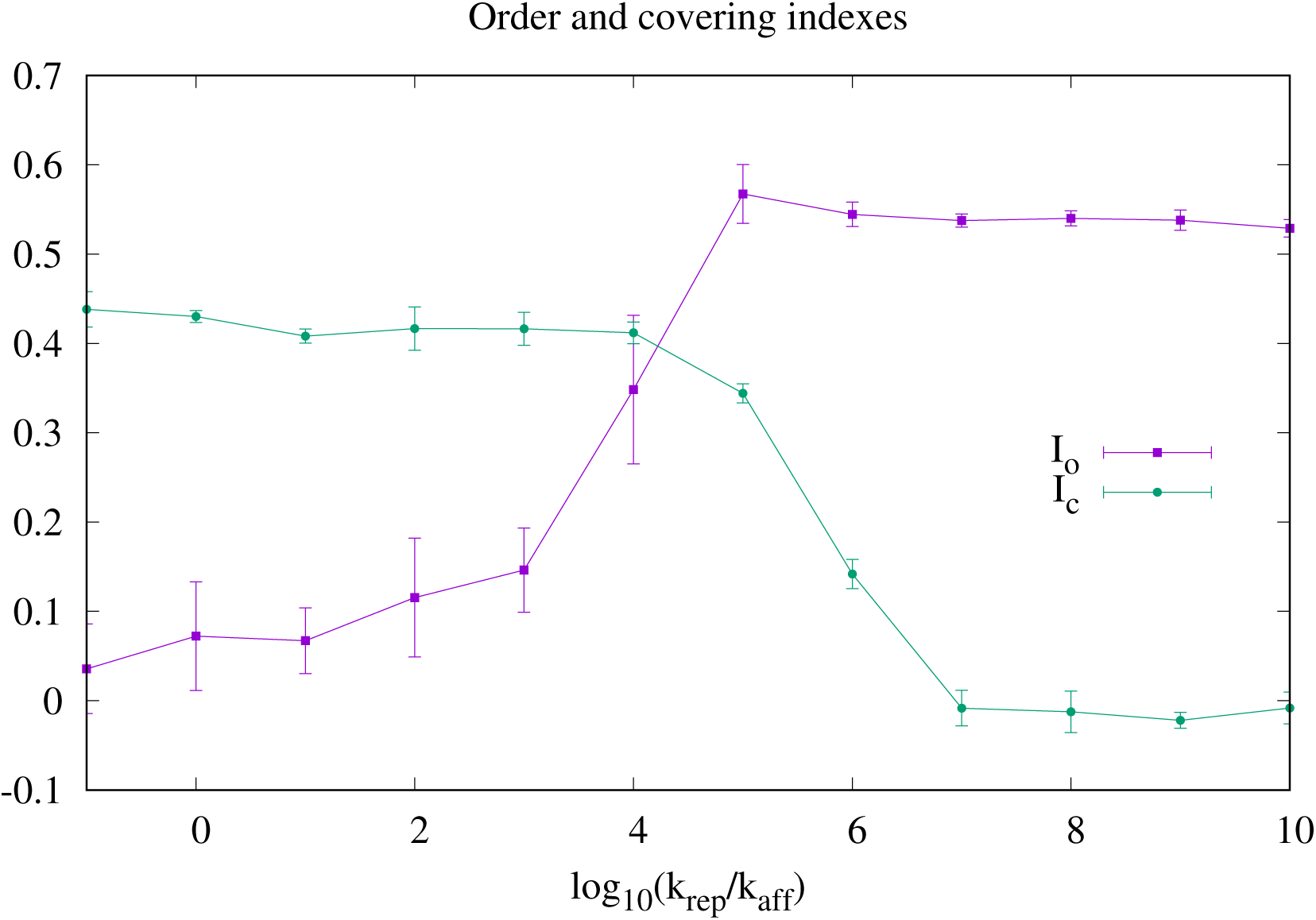

### Final values of order and affinity indices for different simulation conditions

The points represent mean values computed over five simulations and the error-bars represent the standard deviation.

#### Clonal simulations

In clonal simulations, where patches of the epithelium lack either or both repulsion and affinity, the set of clone regions was called *Z*, and the new interaction constants 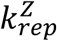 and 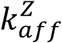. The energies depending on the position of the MCCs (inside or outside *Z*) were thus modified. The repulsion energy reads

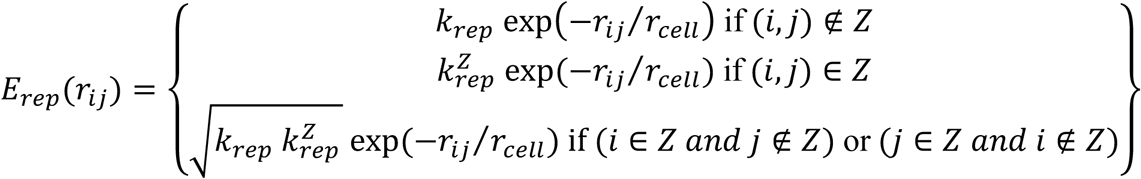

for *r*_*ij*_ ⩾ *r*_*cell*_ (the hard-core energy of *E*_*rep*_(*r*_*ij*_) in section *Energies* is unchanged when *r*_*ij*_ < *r*_*cell*_), and the affinity energy reads 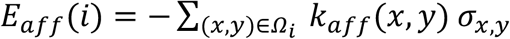 with 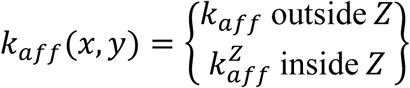

## Movie titles

**Movie 1**: MCCs move actively within the plane of the ectoderm inner layer.

**Movie 2**: Dynamic behaviour of the actin cytoskeleton in immature MCCs.

**Movie 3**: In a simulation dominated by MCC affinity for outer-layer junctions, MCCs home to vertices, but do not disperse properly.

**Movie 4**: In a simulation dominated by MCC mutual repulsion, MCCs disperse properly, but do not home to vertices.

**Movie 5**: In a simulation where MCC mutual repulsion and affinity for outer-layer junctions are balanced, MCCs disperse and home correctly to vertices.

**Movie 6**: When MCCs come into contact via lamellipodia-like structures, they withdraw from each other.

**Movie 7**: MCCs preferentially move along outer-layer intercellular junctions.

**Movie 8**: MCC mutual repulsion is strongly decreased in axitinib-treated embryos.

**Movie 9**: MCC mutual repulsion is strongly decreased in *kit* MO-injected embryos.

**Movie 10**: MCC mutual repulsion is strongly decreased in *scf* MO-injected embryos.

**Movie 11**: In *scf* MO mosaic embryos, MCCs leave Scf-depleted regions without following outer-layer intercellular junctions.

**Movie 12**: Mathematical modelling predicts that MCCs tend to accumulate at boundaries between regions devoid of both repulsion and affinity, and normal regions.

**Movie 13**: When only repulsion is suppressed in clones surrounded by normal regions, MCCs remain within the clones but do not disperse homogeneously.

**Movie 14**: When only affinity is suppressed in clones surrounded by normal regions, MCCs disperse homogeneously within the clones but do not preferentially associate to vertices.

**Movie 15**: MCCs overexpressing Scf-GFP form stable aggregates.

**Movie S1**: Fluorescent labelling of activated caspases 3 and 7 reveals no apoptotic cell death within the ectoderm.

**Movie S2**: Tilted 3D projection from movie 7 showing MCC apically-directed protrusions occurring in correspondence of outer-layer intercellular junctions.

**Movie S3**: MCC mutual repulsion is strongly decreased in CK-666-treated embryos.

**Movie S4**: A GFP-tagged version of ARP2/3 subunit p41 localizes to lamellipodia-like structures and is lost following their retraction upon MCC mutual contact.

**Movie S5**: MCC mutual repulsion is strongly decreased in dn-Kit-injected embryos.

## Notes

### Competing Interest Statement

The authors have declared no competing interest.

## REFERENCES

Belin, B.J., Goins, L.M., and Mullins, R.D. (2014). Comparative analysis of tools for live cell imaging of actin network architecture. Bioarchitecture 4, 189–202.

Benito-Gutierrez, E., Nake, C., Llovera, M., Comella, J.X., and Garcia-Fernandez, J. (2005). The single AmphiTrk receptor highlights increased complexity of neurotrophin signalling in vertebrates and suggests an early role in developing sensory neuroepidermal cells. Development 132, 2191–2202.

Brannan, C.I., Lyman, S.D., Williams, D.E., Eisenman, J., Anderson, D.M., Cosman, D., Bedell, M.A., Jenkins, N.A., and Copeland, N.G. (1991). Steel-Dickie mutation encodes a c-kit ligand lacking transmembrane and cytoplasmic domains. Proc Natl Acad Sci U S A 88, 4671–4674.

Briggs, J.A., Weinreb, C., Wagner, D.E., Megason, S., Peshkin, L., Kirschner, M.W., and Klein, A.M. (2018). The dynamics of gene expression in vertebrate embryogenesis at single-cell resolution. Science 360.

Briscoe, J., and Small, S. (2015). Morphogen rules: design principles of gradient-mediated embryo patterning. Development 142, 3996–4009.

Castillo-Briceno, P., and Kodjabachian, L. (2014). Xenopus embryonic epidermis as a mucociliary cellular ecosystem to assess the effect of sex hormones in a non-reproductive context. Front Zool 11, 9.

Chung, M.I., Kwon, T., Tu, F., Brooks, E.R., Gupta, R., Meyer, M., Baker, J.C., Marcotte, E.M., and Wallingford, J.B. (2014). Coordinated genomic control of ciliogenesis and cell movement by RFX2. Elife 3, e01439.

Cibois, M., Luxardi, G., Chevalier, B., Thome, V., Mercey, O., Zaragosi, L.E., Barbry, P., Pasini, A., Marcet, B., and Kodjabachian, L. (2015). BMP signalling controls the construction of vertebrate mucociliary epithelia. Development 142, 2352–2363.

Cibois, M., Scerbo, P., Thomé, V., Pasini, A. and Kodjabachian, L. (2014). Induction and Differentiation of the Xenopus Ciliated Embryonic Epidermis. In Xenopus Development, M.K.a.J.Z. Kubiak, ed. (Oxford: John Wiley & Sons, Inc).

Davidson, E.H. (2010). Emerging properties of animal gene regulatory networks. Nature 468, 911–920.

Davis, J.R., Huang, C.Y., Zanet, J., Harrison, S., Rosten, E., Cox, S., Soong, D.Y., Dunn, G.A., and Stramer, B.M. (2012). Emergence of embryonic pattern through contact inhibition of locomotion. Development 139, 4555–4560.

Deblandre, G.A., Wettstein, D.A., Koyano-Nakagawa, N., and Kintner, C. (1999). A two-step mechanism generates the spacing pattern of the ciliated cells in the skin of Xenopus embryos. Development 126, 4715–4728.

Flanagan, J.G., Chan, D.C., and Leder, P. (1991). Transmembrane form of the kit ligand growth factor is determined by alternative splicing and is missing in the Sld mutant. Cell 64, 1025–1035.

Fuerst, P.G., Koizumi, A., Masland, R.H., and Burgess, R.W. (2008). Neurite arborization and mosaic spacing in the mouse retina require DSCAM. Nature 451, 470–474.

Galli-Resta, L., Novelli, E., and Viegi, A. (2002). Dynamic microtubule-dependent interactions position homotypic neurones in regular monolayered arrays during retinal development. Development 129, 3803–3814.

Goldman, D.C., Berg, L.K., Heinrich, M.C., and Christian, J.L. (2006). Ectodermally derived steel/stem cell factor functions non-cell autonomously during primitive erythropoiesis in Xenopus. Blood 107, 3114–3121.

Gommerman, J.L., Sittaro, D., Klebasz, N.Z., Williams, D.A., and Berger, S.A. (2000). Differential stimulation of c-Kit mutants by membrane-bound and soluble Steel Factor correlates with leukemic potential. Blood 96, 3734–3742.

Gu, Y., Runyan, C., Shoemaker, A., Surani, A., and Wylie, C. (2009). Steel factor controls primordial germ cell survival and motility from the time of their specification in the allantois, and provides a continuous niche throughout their migration. Development 136, 1295–1303.

Gu, Y., Runyan, C., Shoemaker, A., Surani, M.A., and Wylie, C. (2011). Membrane-bound steel factor maintains a high local concentration for mouse primordial germ cell motility, and defines the region of their migration. PLoS One 6, e25984.

Hayes, J.M., Kim, S.K., Abitua, P.B., Park, T.J., Herrington, E.R., Kitayama, A., Grow, M.W., Ueno, N., and Wallingford, J.B. (2007). Identification of novel ciliogenesis factors using a new in vivo model for mucociliary epithelial development. Dev Biol 312, 115–130.

Heissig, B., Hattori, K., Dias, S., Friedrich, M., Ferris, B., Hackett, N.R., Crystal, R.G., Besmer, P., Lyden, D., Moore, M.A., et al. (2002). Recruitment of stem and progenitor cells from the bone marrow niche requires MMP-9 mediated release of kit-ligand. Cell 109, 625–637.

Hetrick, B., Han, M.S., Helgeson, L.A., and Nolen, B.J. (2013). Small molecules CK-666 and CK-869 inhibit actin-related protein 2/3 complex by blocking an activating conformational change. Chem Biol 20, 701–712.

Higashi, T., Arnold, T.R., Stephenson, R.E., Dinshaw, K.M., and Miller, A.L. (2016). Maintenance of the Epithelial Barrier and Remodeling of Cell-Cell Junctions during Cytokinesis. Curr Biol 26, 1829–1842.

Hu-Lowe, D.D., Zou, H.Y., Grazzini, M.L., Hallin, M.E., Wickman, G.R., Amundson, K., Chen, J.H., Rewolinski, D.A., Yamazaki, S., Wu, E.Y., et al. (2008). Nonclinical antiangiogenesis and antitumor activities of axitinib (AG-013736), an oral, potent, and selective inhibitor of vascular endothelial growth factor receptor tyrosine kinases 1, 2, 3. Clin Cancer Res 14, 7272–7283.

Hultman, K.A., Bahary, N., Zon, L.I., and Johnson, S.L. (2007). Gene Duplication of the zebrafish kit ligand and partitioning of melanocyte development functions to kit ligand a. PLoS Genet 3, e17.

Ing-Esteves, S., Kostadinov, D., Marocha, J., Sing, A.D., Joseph, K.S., Laboulaye, M.A., Sanes, J.R., and Lefebvre, J.L. (2018). Combinatorial Effects of Alpha- and Gamma-Protocadherins on Neuronal Survival and Dendritic Self-Avoidance. J Neurosci 38, 2713–2729.

Irion, U., Frohnhofer, H.G., Krauss, J., Colak Champollion, T., Maischein, H.M., Geiger-Rudolph, S., Weiler, C., and Nusslein-Volhard, C. (2014). Gap junctions composed of connexins 41.8 and 39.4 are essential for colour pattern formation in zebrafish. Elife 3, e05125.

Iwashita, M., Watanabe, M., Ishii, M., Chen, T., Johnson, S.L., Kurachi, Y., Okada, N., and Kondo, S. (2006). Pigment pattern in jaguar/obelix zebrafish is caused by a Kir7.1 mutation: implications for the regulation of melanosome movement. PLoS Genet 2, e197.

Jordan, S.A., and Jackson, I.J. (2000). MGF (KIT ligand) is a chemokinetic factor for melanoblast migration into hair follicles. Dev Biol 225, 424–436.

Kaltenbach, S.L., Yu, J.K., and Holland, N.D. (2009). The origin and migration of the earliest-developing sensory neurons in the peripheral nervous system of amphioxus. Evol Dev 11, 142–151.

Kim, K., Lake, B.B., Haremaki, T., Weinstein, D.C., and Sokol, S.Y. (2012). Rab11 regulates planar polarity and migratory behavior of multiciliated cells in Xenopus embryonic epidermis. Dev Dyn 241, 1385–1395.

Kim, S., Ma, L., Shokhirev, M.N., Quigley, I., and Kintner, C. (2018). Multicilin and activated E2f4 induce multiciliated cell differentiation in primary fibroblasts. Sci Rep 8, 12369.

Lennartsson, J., and Ronnstrand, L. (2012). Stem cell factor receptor/c-Kit: from basic science to clinical implications. Physiol Rev 92, 1619–1649.

Ma, L., Quigley, I., Omran, H., and Kintner, C. (2014). Multicilin drives centriole biogenesis via E2f proteins. Genes Dev 28, 1461–1471.

Mani, M., Venkatasubrahmanyam, S., Sanyal, M., Levy, S., Butte, A., Weinberg, K., and Jahn, T. (2009). Wiskott-Aldrich syndrome protein is an effector of Kit signaling. Blood 114, 2900–2908.

Marchal, L., Luxardi, G., Thome, V., and Kodjabachian, L. (2009). BMP inhibition initiates neural induction via FGF signaling and Zic genes. Proc Natl Acad Sci U S A 106, 17437–17442.

Martin, B.L., and Harland, R.M. (2004). The developmental expression of two Xenopus laevis steel homologues, Xsl-1 and Xsl-2. Gene Expr Patterns 5, 239–243.

Meininger, C.J., Yano, H., Rottapel, R., Bernstein, A., Zsebo, K.M., and Zetter, B.R. (1992). The c-kit receptor ligand functions as a mast cell chemoattractant. Blood 79, 958–963.

Miyazawa, K., Williams, D.A., Gotoh, A., Nishimaki, J., Broxmeyer, H.E., and Toyama, K. (1995). Membrane-bound Steel factor induces more persistent tyrosine kinase activation and longer life span of c-kit gene-encoded protein than its soluble form. Blood 85, 641–649.

Quigley, I.K., Stubbs, J.L., and Kintner, C. (2011). Specification of ion transport cells in the Xenopus larval skin. Development 138, 705–714.

Rizvi, S.A., Neidt, E.M., Cui, J., Feiger, Z., Skau, C.T., Gardel, M.L., Kozmin, S.A., and Kovar, D.R. (2009). Identification and characterization of a small molecule inhibitor of formin-mediated actin assembly. Chem Biol 16, 1158–1168.

Runyan, C., Schaible, K., Molyneaux, K., Wang, Z., Levin, L., and Wylie, C. (2006). Steel factor controls midline cell death of primordial germ cells and is essential for their normal proliferation and migration. Development 133, 4861–4869.

Samayawardhena, L.A., Kapur, R., and Craig, A.W. (2007). Involvement of Fyn kinase in Kit and integrin-mediated Rac activation, cytoskeletal reorganisation, and chemotaxis of mast cells. Blood 109, 3679–3686.

Sedzinski, J., Hannezo, E., Tu, F., Biro, M., and Wallingford, J.B. (2016). Emergence of an Apical Epithelial Cell Surface In Vivo. Dev Cell 36, 24–35.

Sedzinski, J., Hannezo, E., Tu, F., Biro, M., and Wallingford, J.B. (2017). RhoA regulates actin network dynamics during apical surface emergence in multiciliated epithelial cells. J Cell Sci 130, 420–428.

Sirour, C., Hidalgo, M., Bello, V., Buisson, N., Darribere, T., and Moreau, N. (2011). Dystroglycan is involved in skin morphogenesis downstream of the Notch signaling pathway. Mol Biol Cell 22, 2957–2969.

Stubbs, J.L., Davidson, L., Keller, R., and Kintner, C. (2006). Radial intercalation of ciliated cells during Xenopus skin development. Development 133, 2507–2515.

Stubbs, J.L., Vladar, E.K., Axelrod, J.D., and Kintner, C. (2012). Multicilin promotes centriole assembly and ciliogenesis during multiciliate cell differentiation. Nat Cell Biol 14, 140–147.

Suraneni, P., Fogelson, B., Rubinstein, B., Noguera, P., Volkmann, N., Hanein, D., Mogilner, A., and Li, R. (2015). A mechanism of leading-edge protrusion in the absence of Arp2/3 complex. Mol Biol Cell 26, 901–912.

Suraneni, P., Rubinstein, B., Unruh, J.R., Durnin, M., Hanein, D., and Li, R. (2012). The Arp2/3 complex is required for lamellipodia extension and directional fibroblast cell migration. J Cell Biol 197, 239–251.

Tabone-Eglinger, S., Calderin-Sollet, Z., Pinon, P., Aebischer, N., Wehrle-Haller, M., Jacquier, M.C., Boettiger, D., and Wehrle-Haller, B. (2014). Niche anchorage and signaling through membrane-bound Kit-ligand/c-kit receptor are kinase independent and imatinib insensitive. FASEB J 28, 4441–4456.

Tabone-Eglinger, S., Wehrle-Haller, M., Aebischer, N., Jacquier, M.C., and Wehrle-Haller, B. (2012). Membrane-bound Kit ligand regulates melanocyte adhesion and survival, providing physical interaction with an intraepithelial niche. FASEB J 26, 3738–3753.

Tajima, Y., Moore, M.A., Soares, V., Ono, M., Kissel, H., and Besmer, P. (1998). Consequences of exclusive expression in vivo of Kit-ligand lacking the major proteolytic cleavage site. Proc Natl Acad Sci U S A 95, 11903–11908.

Trindade, M., Messenger, N., Papin, C., Grimmer, D., Fairclough, L., Tada, M., and Smith, J.C. (2003). Regulation of apoptosis in theXenopus embryo by Bix3. Development 130, 4611–4622.

Villar-Cervino, V., Molano-Mazon, M., Catchpole, T., Valdeolmillos, M., Henkemeyer, M., Martinez, L.M., Borrell, V., and Marin, O. (2013). Contact repulsion controls the dispersion and final distribution of Cajal-Retzius cells. Neuron 77, 457–471.

Walderich, B., Singh, A.P., Mahalwar, P., and Nusslein-Volhard, C. (2016). Homotypic cell competition regulates proliferation and tiling of zebrafish pigment cells during colour pattern formation. Nat Commun 7, 11462.

Wehrle-Haller, B., Meller, M., and Weston, J.A. (2001). Analysis of melanocyte precursors in Nf1 mutants reveals that MGF/KIT signaling promotes directed cell migration independent of its function in cell survival. Dev Biol 232, 471–483.

Wehrle-Haller, B., and Weston, J.A. (1995). Soluble and cell-bound forms of steel factor activity play distinct roles in melanocyte precursor dispersal and survival on the lateral neural crest migration pathway. Development 121, 731–742.

Werner, M.E., and Mitchell, B.J. (2012). Understanding ciliated epithelia: the power of Xenopus. Genesis 50, 176–185.

Werner, M.E., Mitchell, J.W., Putzbach, W., Bacon, E., Kim, S.K., and Mitchell, B.J. (2014). Radial intercalation is regulated by the Par complex and the microtubule-stabilizing protein CLAMP/Spef1. J Cell Biol 206, 367–376.

Zuo, K., Kuang, D., Wang, Y., Xia, Y., Tong, W., Wang, X., Chen, Y., Duan, Y., and Wang, G. (2016). SCF/c-kit transactivates CXCR4-serine 339 phosphorylation through G protein-coupled receptor kinase 6 and regulates cardiac stem cell migration. Sci Rep 6, 26812.

