## Supplementary material for "The SCF/KIT pathway implements self-organised epithelial patterning by cell movement": Supp figures and table

### SUPPLEMENTARY FIGURES

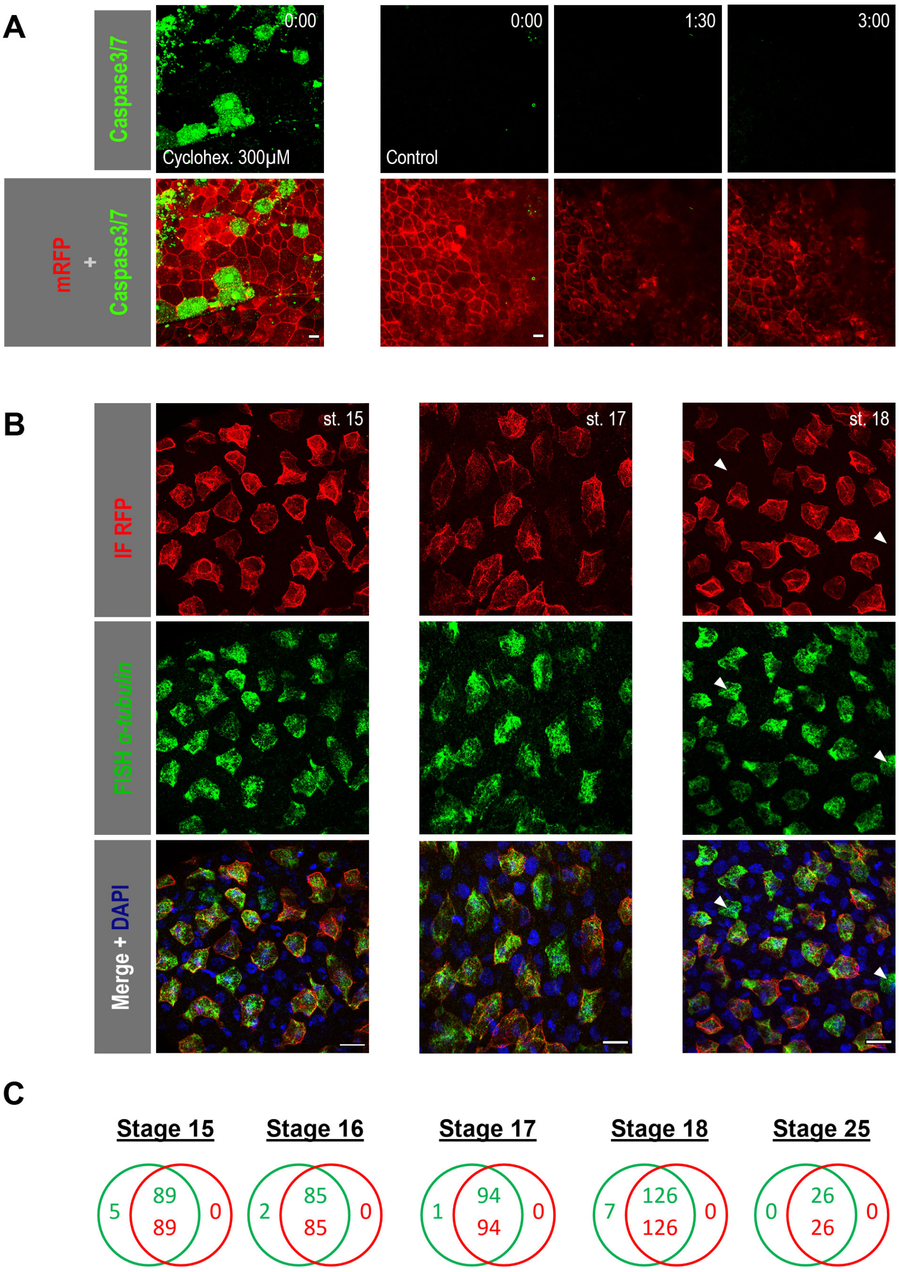


**Figure S1**

**Figure S1. Cell death and cell fate reversal are unlikely to play a major role in establishing the MCC regular pattern.**

**A:** Stage 15 mRFP-injected *Xenopus* embryos were incubated with the CellEvent™ Caspase-3/7 Green Detection Reagent to detect activation of the executioner Caspases 3 and 7 in presence of 300µM cycloheximide as an apoptosis inducer (Trindade et al., Development, 2003) (left) or of 1% (v/v) DMSO as a control and filmed for 3h (right). Apoptotic cells (green) were found in the epidermis of cycloheximide-treated embryos (left), but not in control ones, even after prolonged observation (right). Embryos were injected with mRFP (red) to allow a better visualisation of cell outlines. Scale bar is 20µm. **B:** α-tub::RFP transgenic embryos were fixed at different developmental stages and subjected to IF with an anti-RFP antibody (red, top row) and FISH against *α-tubulin* (green, middle row). Merging the two channels (bottom row) showed few *α-tub*-positive, RFP-negative cells (white arrowheads), but failed to uncover any RFP-positive, *α-tub*-negative cells. DAPI staining (blue, bottom row) allowed the identification of nuclei. Scale bar is 20µm. **C:** Venn diagrams showing the overlap of the GFP- and RFP-positive cell populations at different developmental stages.

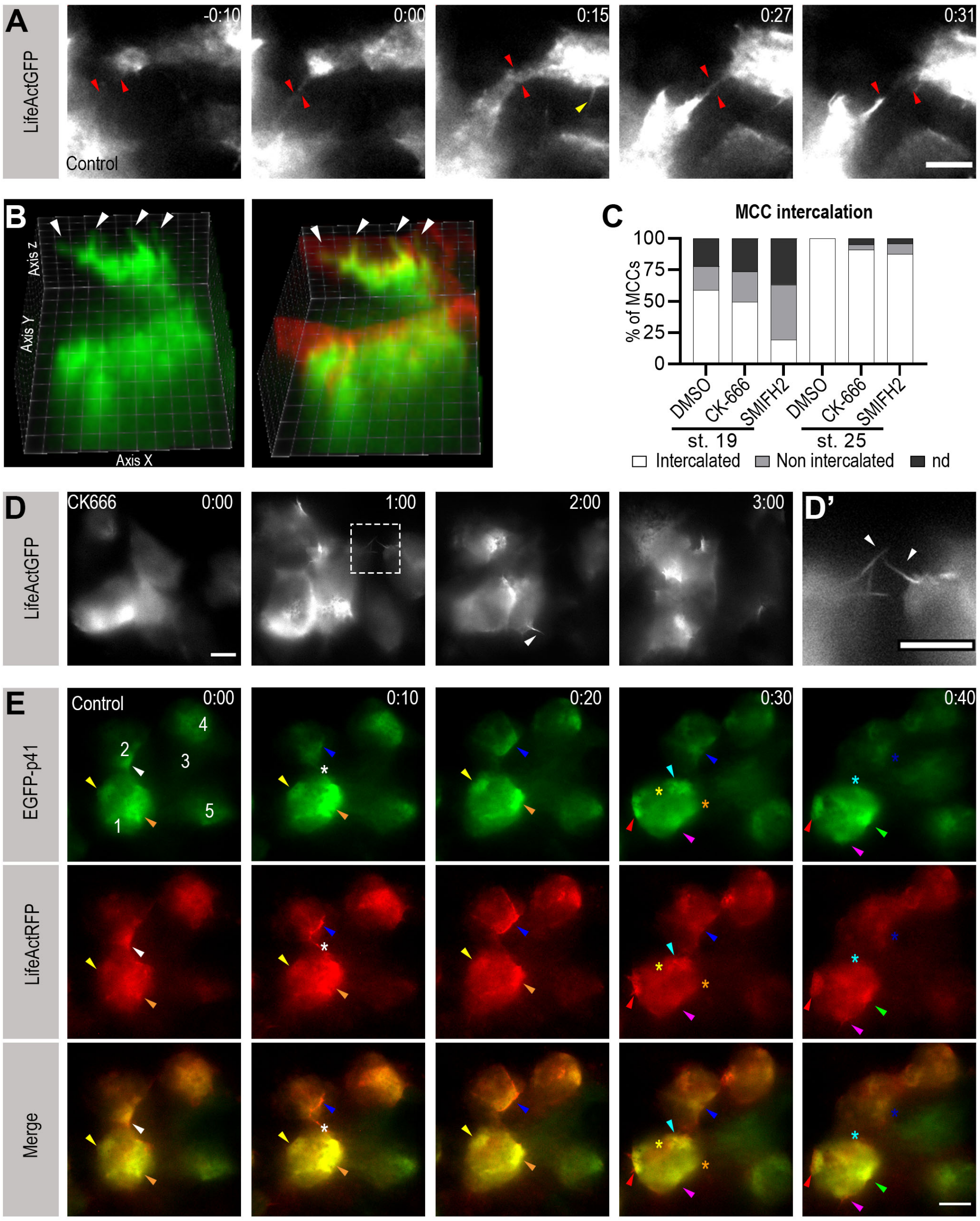


Figure S2

**Figure S2. Role of actin cytoskeleton in MCC patterning.**

**A:** Frames from a movie showing LifeActGFP-expressing MCCs establishing transient contact via lamellipodia-like structures (red arrowheads) that were subsequently retracted. The yellow arrowhead points to an example of thinner, filopodia-like structures. Scale bar is 20µm. **B:** A tilted 3D projection from movie S2 showing that apically directed protrusions (white arrowheads) corresponded to outer-layer cell junctions. **C:** Quantification of MCC intercalation at stage 19 and stage 25 in control embryos and embryos treated with CK-666 or SMIFH2. **D, D':** Frames from movie S3 showing that LifeActGFP-expressing MCCs displayed decreased mutual repulsion in CK-666 treated embryos. White arrowheads point to filopodia-like structures. Scale bar is 20µm. **E:** Frames from movie S4 showing that local accumulations of a GFP-tagged version of the branched actin-nucleator ARP2/3 subunit p41 colocalised with the LifeActRFP signal in lamellipodia-like structures (arrowheads), but were lost following MCC mutual contact (asterisks). White numbers show different MCCs. Scale bar is 20µm.

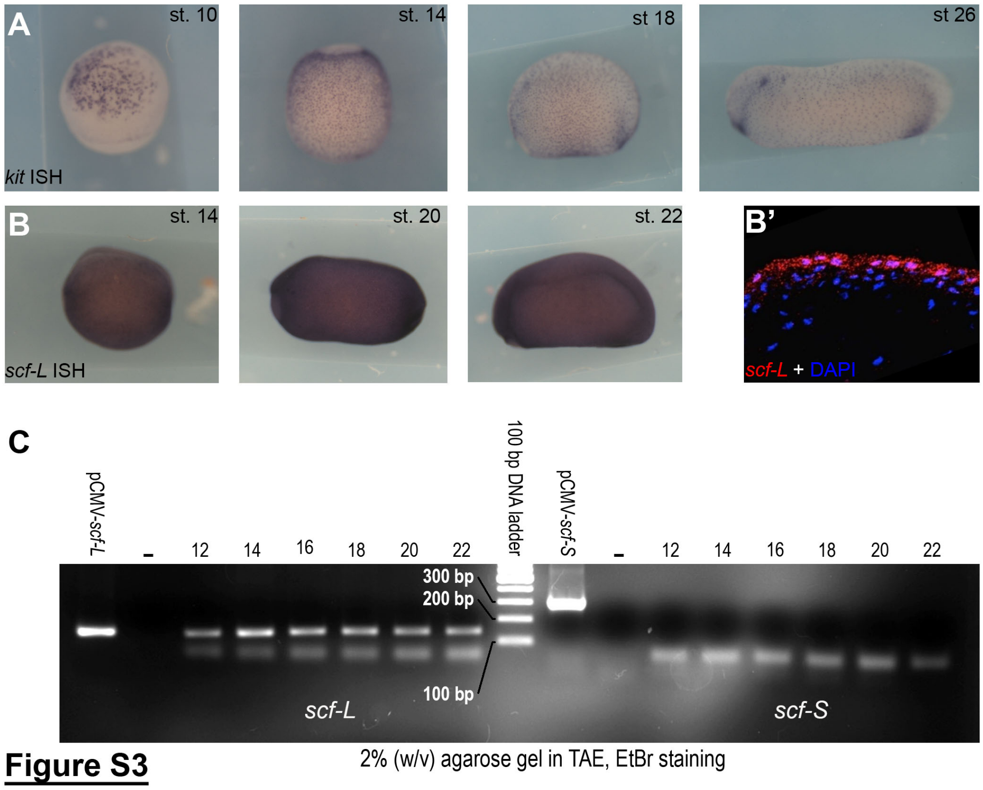


**Figure S3. Expression of *kit* and *scf* in *Xenopus* epidermis.**
**A:** Whole-mount ISH with a probe against *kit-L* and *kit-S* revealed expression in scattered non-neural ectoderm cells at stages 12 to 24. Anterior is to the top for stages 12 and 14, to the left for stages 18 and 24. **B:** Whole-mount ISH with a probe against *scf-L* revealed expression throughout the ectoderm from stages 14 to 22. Anterior to the left. **B’:** FISH on a sectioned embryo at stage 25 revealed stronger *scf-L* expression (red) in the ectoderm outer layer. DAPI (blue) stained the nuclei. **C:** RT-PCR on total RNA from animal caps at different developmental stages with primers encompassing the exon6 region of *scf-L* (left) or *scf-S* (right) amplified a 141bp band, only for *scf-L*, which was present at all developmental stages analysed. Plasmids containing the full-length cDNA of either *scf-L* (pCMV-Sport6-Scf-L) or *scf-S* (pCMV-Sport6-Scf-S) were used as positive controls for amplification. Molecular weight marker is GeneRuler 100bp Plus DNA Ladder (Thermo Scientific).

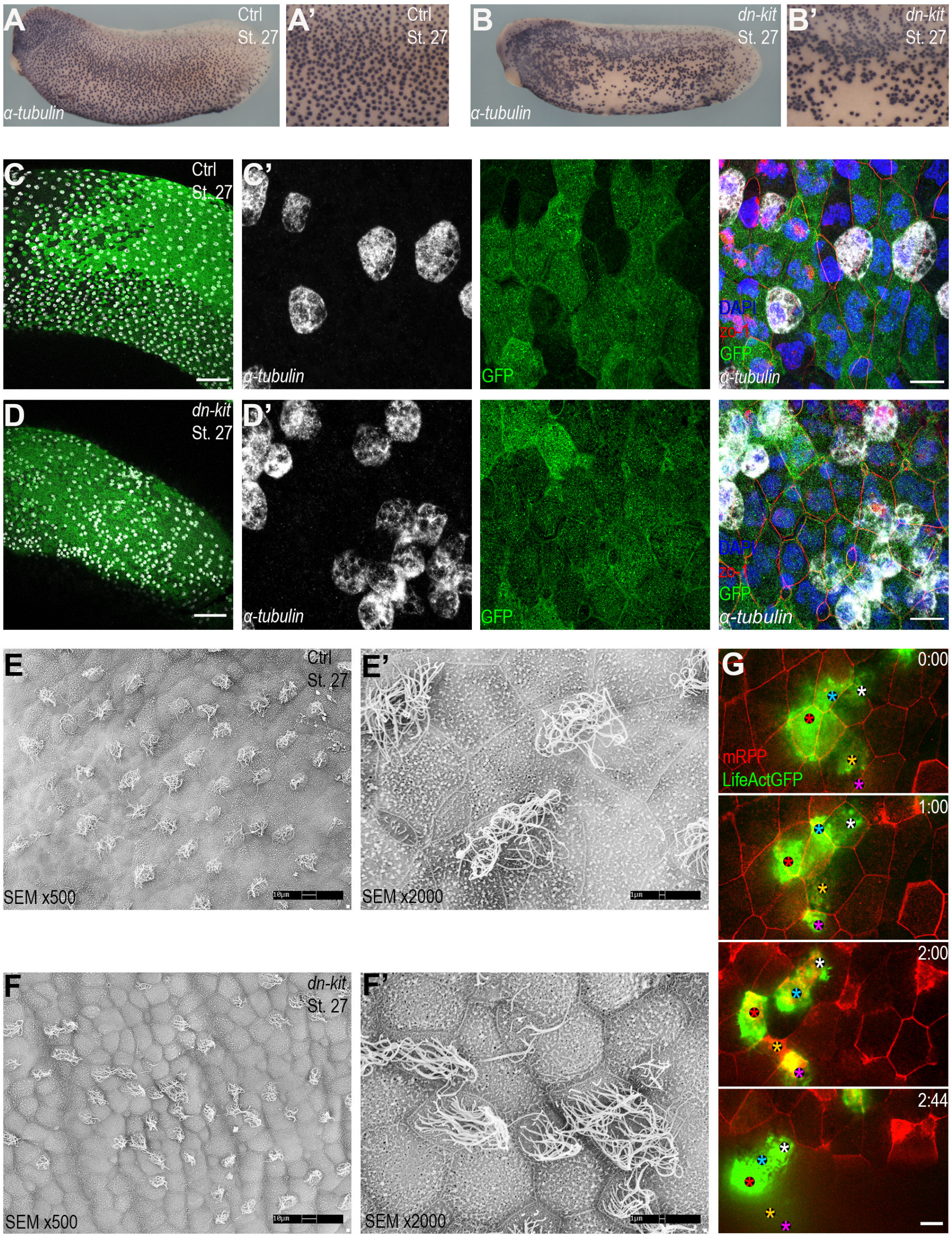


**Figue S4**

**Figure S4. Expression of dominant-negative Kit affects MCC distribution.**

**A, B:** Whole-mount ISH with a probe against *α-tubulin* revealed normal MCC distribution in stage 27 control embryos (A) and patterning irregularities in age-matched *dn-kit* injected embryos (B). **A'** and **B'** are close-ups of A and B respectively. **C, D:** Whole-mount FISH with a probe against *α-tubulin* (white) in control (C) and *dn-kit* injected (D) stage 27 embryos. mRNA coding for GFP (green) was used as a tracer for *dn-kit* mRNA, DAPI (blue) stained nuclei. Scale bar is 200µm. **C'** and **D'** are close-ups of C and D, respectively, showing the irregular clustering of MCCs in *dn-kit* injected embryos. Scale bar is 20µm. **E, F:** scanning electron microscopy images of control (E) and *dn-kit* injected (F) stage 27 embryos. **E'** and **D'** are close-ups of E and D, respectively, showing that MCCs in *dn-kit* injected embryos were irregularly clustered but did produce cilia. **G:** Frames from movie S5, showing that LifeActGFP-positive MCCs failed to separate properly in *dn-kit* injected embryos. mRNA coding for RFP (red) served as tracer for *dn-kit* mRNA. Scale bar is 20µm.

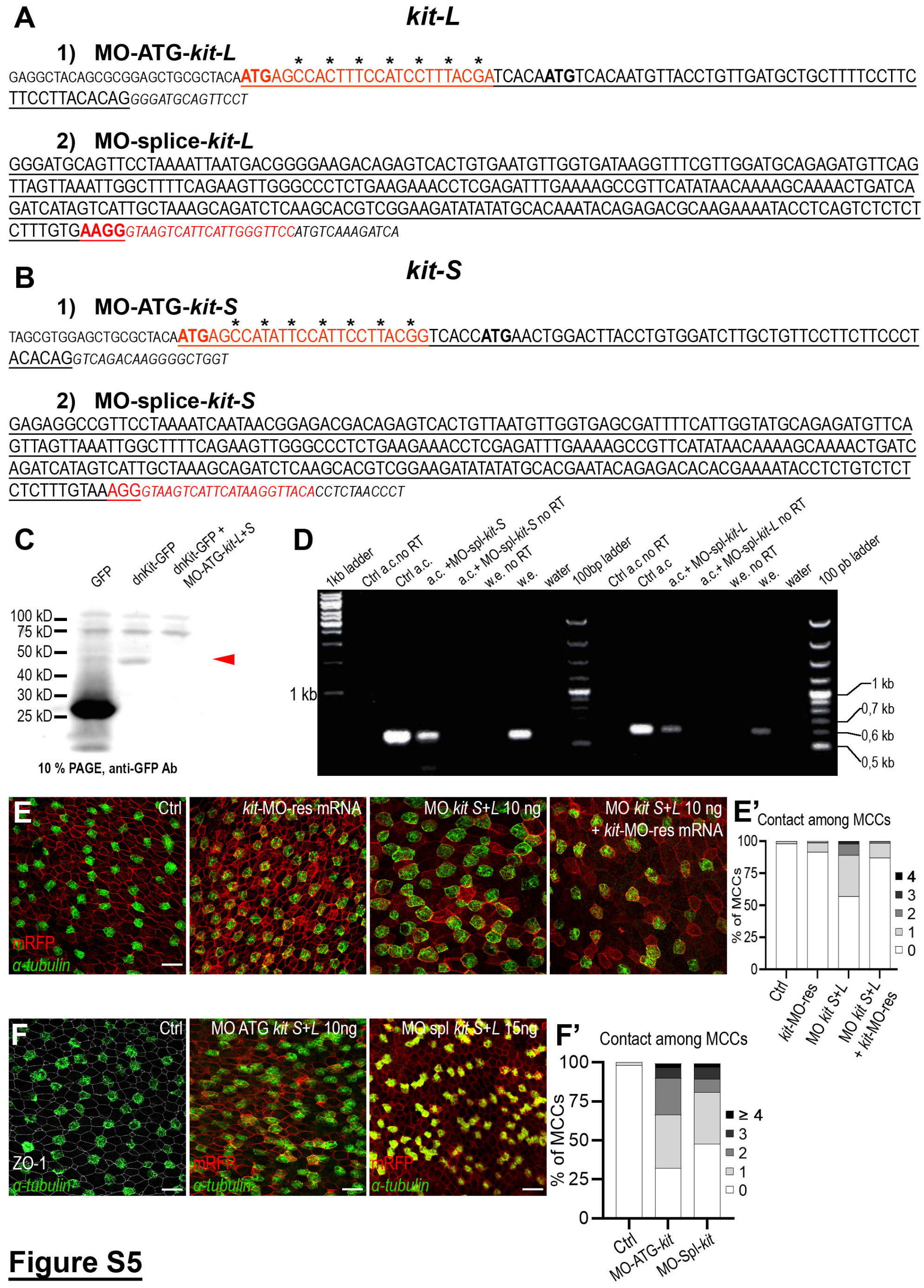


**Figure S5**

**Figure S5. Characterisation of Morpholino Oligos against *kit-L* and *kit-S*.**
**A:** Morpholino Oligos against *kit-L*. **1)** Upper case underlined: *kit-L* exon 1; lower case: 5'UTR; lower case italic: intron 1; in red the stretch targeted by MO-ATG-kit-L. **2)** Upper case underlined: *kit-L* exon 2; lower case italic: intron 2; in red the stretch targeted by MO-splice-kit-L. **B:** Morpholino Oligos against *kit-S*. **1)** Upper case underlined: *kit-S* exon 1; lower case: 5'UTR; lower case italic: intron 1; in red the stretch targeted by MO-ATG-kit-S. The asterisk/arrowheads point to the seven bases mutated to generate MOres-kit-S. **2)** Upper case underlined: *kit-S* exon 2; lower case italic: intron 2; in red the stretch targeted by MO-splice-kit-S. **C:** Lysates from animal caps injected with mRNA coding for GFP and mRNA coding for a C-terminally GFP-tagged dominant-negative form of Kit-S alone or together with MO-

ATG-kit-L+S were subjected to Western blotting with an anti-GFP antibody. The red arrowhead points to a GFP-positive band corresponding to the intracellular fragment generated by proteolytic cleavage of the dn-Kit-GFP protein, which was lost in embryos co-injected with MO-ATG-kit-L+S. **D:** RT-PCR on *kit-L* and *kit-S* to test the action of the splice-blocking MOs; a.c.: animal caps; w.e.: whole embryos; 0.8% agarose/TAE gel. The injection of MO-splice-kit-S or MO-splice-kit-L resulted in a decrease of the levels of *kit-S* and *kit-L* mRNAs, respectively. **E:** *α-tub* FISH (green) and RFP IF (red) on stage 25 embryos showed, from left to right, normal MCC distribution in control embryos and in embryos injected with mRNA coding for morpholino-resistant form MOres-kit-S alone, abnormal MCC clustering in embryos injected with MO-ATG-kit-L+S, and recovery of normal MCC distribution in embryos co-injected with MO-ATG-kit-L+S and MOres-kit-*S* mRNA. In all cases, mRNA for RFP was used as an injection tracer. Scale bar is 20μm. **E':** A quantification of MCC clustering at stage 25 in control embryos, embryos injected with MOres-kit-S mRNA alone, MO-ATG-kit-L+S alone and MO-ATG-kit-L+S together with MOres-kit-S mRNA. **F:** *α-tub* FISH (green) and ZO-1 IF (white) on control embryos (left panel) and embryos injected with either MO-ATG-kit-L+S (middle panel) or MO-splice-kit-L+S (right panel) showed that both ATG- and splice-targeting MOs led to the same phenotype of irregularly distributed and clustered MCCs at stage 25. mRFP mRNA (red) was used as MO tracer. Scale bar is 20μm. The graph in **F'** shows a quantification of the abnormal MCC clustering in MO-ATG-kit-L+S and MO-splice-kit-L+S injected embryos.

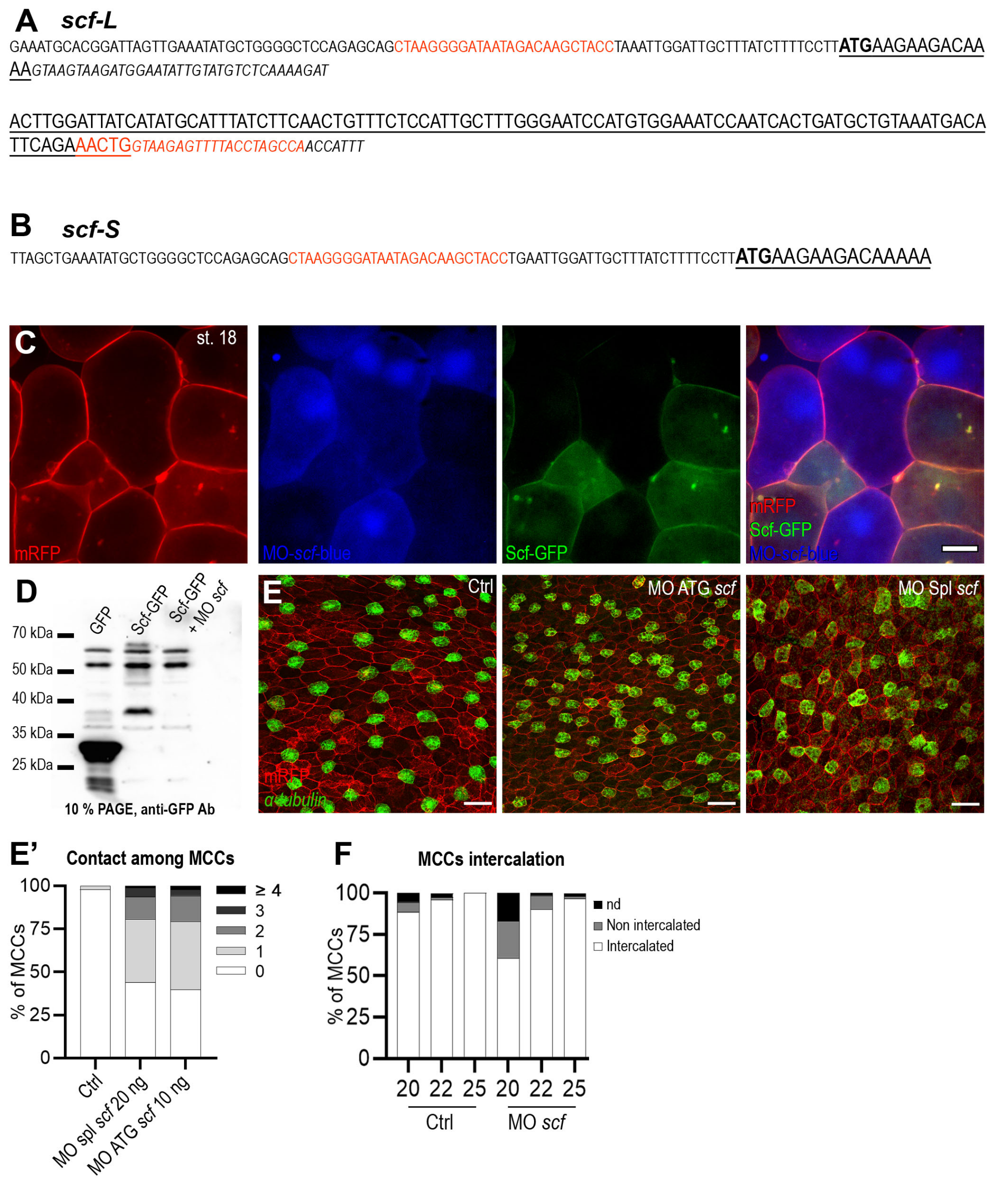


### Figure S6

**Figure S6. Characterisation of Morpholino Oligos against *scf-L* and *scf-S*.**
**A:** Morpholino Oligos against *scf-L*. **1)** Upper case underlined: *scf-L* exon 1; lower case: 5'UTR; lower case italic: intron 1; in red the stretch targeted by MO-ATG-scf-L/S. **2)** Upper case underlined: *scf-L* exon 2; lower case italic: intron 2; in red the stretch targeted by MO-splice-scf-L. **B:** Morpholino Oligos against *scf-S*. **1)** Upper case underlined: *scf-S* exon 1; lower case: 5'UTR; in red the stretch targeted by MO-ATG-scf-L/S. **C:** Top view of live outer-layer cells from a stage 18 embryo co-injected with mRNAs coding for Scf-GFP (green) and RFP (red), as well as a blue-tagged version of MO-ATG-scf-L/S. The Scf-GFP signal was lost from the MO-receiving cells, while the RFP signal was unchanged. Scale bar is 20µm. **D:** Lysates from animal caps injected with mRNA coding for GFP, Scf-GFP and Scf-GFP together with MO-ATG-scf-L/S were subjected to Western blotting with an anti-GFP antibody. Bands corresponding to the 55kDa full-length Scf-GFP and to the 37kDa exon 6 proteolytic cleavage product were absent from MO-ATG-scf-L/S injected embryos. **E:** *α-tub* FISH (green) and RFP (red) IF on control embryos (left panel) and embryos injected with either MO-ATG-scf-L/S (middle panel) or MO-splice-scf-L (right panel) showed that both ATG- and splice-targeting

MOs led to the same phenotype of irregularly distributed and clustered MCCs at stage 25. mRFP mRNA (red) was used as MO tracer. Scale bar is 20µm. **E'** shows a quantification of the abnormal MCC clustering in MO-splice-scf-L/S and MO-ATG-scf-L/S-injected embryos. **F**: A quantification of the MCC intercalation in control and *scf* MO-injected embryos.

| Figure | Stage/ condition | n cells | n embryos | n experiments |
|---|---|---|---|---|
| 1 B | 14 | 456 | 4 | 3 |
| | 16 | 591 | 5 | 3 |
| | 18 | 408 | 4 | 3 |
| | 20 | 761 | 6 | 3 |
| | 22 | 572 | 6 | 3 |
| | 25 | 450 | 6 | 3 |
| 1 C | 14 | 422 | 3 | 2 |
| | 16 | 587 | 4 | 2 |
| | 18 | 306 | 3 | 2 |
| | 20 | 331 | 2 | 1 |
| 1 D | 14 | 440 | 3 | 2 |
| | 16 | 490 | 4 | 3 |
| | 18 | 306 | 3 | 2 |
| | 20 | 331 | 2 | 1 |
| | 22 | 384 | 4 | 2 |
| | 25 | 227 | 4 | 2 |
| 3 F | Ctrl | 108 | 4 | 4 |
| | CK-666 | 62 | 5 | 5 |
| 3 G | Ctrl | 94 | 10 | 10 |
| | CK-666 | 26 | 3 | 3 |
| 3H | Ctrl | 227 | 4 | 4 |
| | CK-666 | 209 | 2 | 2 |
| | SMIFH2 | 144 | 2 | 2 |
| 3I | Ctrl | 248 | 4 | 1 |
| | CK-666 | 491 | 6 | 1 |
| | SMIFH2 | 457 | 6 | 1 |
| 5 C | Ctrl | 108 | 4 | 4 |
| | Axtitinib | 64 | 3 | 3 |
| | MO *kit* | 147 | 5 | 5 |
| 5 C' | Ctrl | 94 | 10 | 10 |
| | Axtitinib | 56 | 9 | 9 |
| | MO *kit* | 70 | 5 | 5 |
| 5 G | Ctrl st 20 | 683 | 6 | 2 |
| | Ctrl st 22 | 267 | 4 | 2 |
| | Ctrl st 25 | 377 | 6 | 4 |
| | Axtitinib st 20 | 441 | 5 | 2 |
| | Axtitinib st 22 | 625 | 4 | 2 |
| | Axtitinib st 25 | 724 | 6 | 3 |
| | MO *kit* st 20 | 248 | 2 | 1 |
| | MO *kit* st 22 | 370 | 3 | 1 |
| | MO *kit* st 25 | 411 | 4 | 2 |
| 5 H | Ctrl st 20 | 761 | 6 | 3 |
| | Ctrl st 22 | 572 | 6 | 3 |
| | Ctrl st 25 | 663 | 12 | 4 |
| | Axtitinib st 20 | 830 | 8 | 3 |
| | Axtitinib st 22 | 753 | 5 | 2 |
| | Axtitinib st 25 | 735 | 8 | 2 |
| | MO *kit* st 20 | 860 | 5 | 2 |
| | MO *kit* st 22 | 998 | 8 | 2 |
| | MO *kit* st 25 | 857 | 9 | 3 |

| Figure | Stage/ condition | n cells | n embryos | n experiments |
|---|---|---|---|---|
| 5 I | Ctrl | 227 | 4 | 4 |
| | Axtitinib | 231 | 2 | 2 |
| | MO *kit* | 219 | 2 | 2 |
| 6 C | Ctrl st 20 | 761 | 6 | 3 |
| | Ctrl st 22 | 572 | 6 | 3 |
| | Ctrl st 25 | 518 | 8 | 4 |
| | MO *scf* st 20 | 678 | 7 | 4 |
| | MO *scf* st 22 | 545 | 6 | 1 |
| | MO *scf* st 25 | 768 | 8 | 4 |
| 6 D | Ctrl | 227 | 4 | 4 |
| | MO *scf* | 220 | 3 | 3 |
| 6 F | Ctrl | 108 | 4 | 4 |
| | MO *scf* | 64 | 5 | 5 |
| 6 I | Ctrl | 94 | 10 | 10 |
| | MO *scf* | 62 | 5 | 5 |
| 7 C | Ctrl | 506 | 8 | 2 |
| | paTub::Scf-GFP | 515 | 8 | 2 |
| 7 H | Ctrl | 1707 | 16 | 2 |
| | *kit* mRNA 0,1 ng | 1526 | 15 | 2 |
| | *kit* mRNA 1 ng | 1704 | 17 | 2 |
| 7 I | Ctrl | 272 | 5 | 2 |
| | *kit* mRNA 0,1 ng | 301 | 5 | 2 |
| | *kit* mRNA 1 ng | 299 | 5 | 2 |
| S2 C | Ctrl st 19 | 373 | 3 | 1 |
| | CK-666 st 19 | 288 | 3 | 1 |
| | SMIFH2 st 19 | 386 | 3 | 1 |
| | Ctrl st 25 | 116 | 2 | 1 |
| | CK-666 st 25 | 208 | 2 | 1 |
| | SMIFH2 st 25 | 144 | 2 | 1 |
| S5 E' | Ctrl | 954 | 14 | 2 |
| | *kit*-MO-Res | 837 | 13 | 2 |
| | MO *kit* | 1049 | 14 | 2 |
| | MO *kit* + *kit*-MO-res | 894 | 14 | 2 |
| S5 F' | Ctrl | 558 | 8 | 2 |
| | MO *kit* ATG | 522 | 5 | 2 |
| | MO *kit* spl | 592 | 6 | 2 |
| S6 E' | Ctrl | 602 | 12 | 2 |
| | MO Spl *scf* 20 ng | 624 | 10 | 2 |
| | MO ATG *scf* 10ng | 723 | 7 | 2 |
| S6 F | Ctrl st 20 | 683 | 6 | 2 |
| | Ctrl st 22 | 267 | 4 | 2 |
| | Ctrl st 25 | 226 | 4 | 2 |
| | MO *scf* st 20 | 720 | 7 | 3 |
| | MO *scf* st 22 | 408 | 4 | 1 |
| | MO *scf* st 25 | 367 | 5 | 2 |

**Table S1:** Total number of cells, embryos and repeat experiments for all graphs in all figures
